# A dsRNA Virus Converts Early Endosomes into Stable Replication Organelles Through Maturation Arrest and Retrograde Transport

**DOI:** 10.64898/2026.10.02.756243

**Authors:** Maria Cecilia Gimenez, Diego Ferrero, Mariam Issa, Serene Moussaoui, Micaela Cabrini, Amanda Rebouças Paixão, Liyuan Liu, Suriakarthiga Ganesan, Durga Acharya, Mandy Boermel, Vanina Zaremberg, Nuria Verdaguer, Mauricio Roberto Terebiznik

## Abstract

Infectious bursal disease virus (IBDV) and infectious pancreatic necrosis virus (IPNV) are major animal pathogens that impose a heavy burden on the poultry and aquaculture industries. Yet, major aspects of their unique replication program remain unknown. IBDV is unusual among double-stranded RNA viruses in that it replicates in association with host endosomes. We previously demonstrated this association to be orchestrated by the viral protein VP3 and the early endosomal lipid phosphatidylinositol 3-phosphate [PtdIns(3)P], suggesting an essential role for VP3 in forming birnaviruses’ specialized replication compartments. Here, combining high-resolution fluorescence and correlative light-electron microscopy with diverse molecular and biochemical analyses, we define how the IBDV protein VP3 targets and remodels host endosomes, and establish that endosome-associated replication is conserved in members of the *Birnaviridae* family. We show that IBDV VP3 targets early endosomes via the PtdIns(3)P effector Early Endosomal Antigen 1 (EEA1) to arrest their maturation and prompt their association with the Golgi complex via ESCPE-1-mediated retrograde transport. We further show that IPNV also exploits endosomal membranes for replication in a PtdIns(3)P dependent manner, although for this virus VP3 engages endosomes through an EEA1-independent mechanism yet to be determined. This work uncovers how birnaviruses exploit a single viral protein to orchestrate endosome remodeling and intracellular trafficking to establish specialized replication compartments while revealing both conserved and host-adapted features of birnaviruses replication.

**Significance Statement:** Infectious Bursal Disease Virus (IBDV) and Infectious Pancreatic Necrosis Virus (IPNV) are globally important pathogens that threaten poultry and aquaculture production, causing major economic losses and impacting food security. Birnaviruses are unusual among double-stranded RNA viruses and have been proposed as evolutionary intermediates in RNA virus evolution, yet many aspects of their replication strategy remain unclear. Here, we show that two evolutionarily distinct birnaviruses depend on PtdIns(3)P-positive endosomes for replication and identify VP3 as the viral factor that targets and remodels these compartments. We further define how VP3 redirects early endosomes, promoting their retrograde trafficking toward the Golgi complex and revealing how birnaviruses convert transient endosomal membranes into stable replication platforms.

## Introduction

The *Birnaviridae* family comprises a small group of non-enveloped, icosahedral viruses with a bi-segmented double-stranded RNA (dsRNA) genome packaged within a single-layered T=13 capsid (1). Among its members, Infectious Bursal Disease Virus (IBDV) and Infectious Pancreatic Necrosis Virus (IPNV) are the prototypes of the genera *Avibirnavirus* and *Aquabirnavirus*, respectively. Both are major animal pathogens that impose a substantial burden on the aquaculture and poultry industry (2).

Birnaviruses depart from the canonical architecture of dsRNA viruses by lacking the T=1 inner core, which functions as a transcriptionally active molecular machine throughout the replication cycle, supporting genome replication and shielding viral dsRNA from host cell surveillance (3). Instead, birnaviruses package their genomes into transcriptionally active ribonucleoprotein complexes (RNPs) composed of the dsRNA genome, the viral RNA-dependent RNA polymerase VP1, and the structural protein VP3 (4, 5). VP3 is a multifunctional protein that protects the dsRNA genome, scaffolds capsid assembly, recruits and regulates VP1, modulates innate antiviral responses, and associates with host membranes, a function essential for efficient replication (3, 6). The archetypal birnavirus IBDV employs a replication strategy that more closely resembles those of positive-sense single-stranded RNA viruses (+ssRNA viruses) than those of other dsRNA viruses. In IBDV, RNPs are targeted after virus entry and uncoating to the surface of host endosomes, which are subsequently hijacked as platforms for viral replication (7–10). These membrane-associated replication platforms are analogous to the replication organelles (ROs) assembled by +ssRNA viruses, which provide a compartmentalized environment that supports viral replication while shielding viral genomes and replication intermediates from host antiviral surveillance (11). IBDV-hijacked endosomes accumulate in the perinuclear region, in close association with the Golgi complex, and mounting evidence indicates that they represent specialized organelles with hybrid endosomal characteristics (8, 9, 12, 13).

Consistent with the central role of endosomal membranes in IBDV replication, we previously identified VP3 as the viral factor responsible for targeting RNPs to the surface of host endosomes. This process is mediated by the VP3 PATCH2 domain and depends on phosphatidylinositol 3-phosphate [PtdIns(3)P], a signature phosphoinositide of early endosomal membranes (8, 12). PtdIns(3)P is a key regulator of early endosomal trafficking and maturation, and its accumulation on endosomal membranes is typically transient, as it is converted into other phosphoinositide species as the endosome progress through the sorting and maturation pathway (14). This temporal restriction raises the question of how IBDV maintains stable association of its replication machinery with the highly dynamic endosomal membranes.

Here, we investigated the role of VP3 in endosomal hijacking and identified key host determinants exploited by VP3 to drive organelle remodeling and intracellular trafficking. We show that IBDV VP3 interacts with the PtdIns(3)P effector Early Endosomal Antigen 1 (EEA1) and promotes the persistence of PtdIns(3)P on hijacked endosomes. This interaction preserves the early endosomal state of these compartments (hereinafter referred to as VP3-Es), preventing their progression towards lysosomal degradation while facilitating their association with the Golgi complex through ESCPE-1-dependent retrograde transport. Disruption of this pathway markedly impairs viral infectivity. Furthermore, we demonstrate that the fish pathogen IPNV also exploits endosomal membranes for replication in a PtdIns(3)P-dependent manner. However, unlike IBDV VP3, the IPNV VP3 protein associates with host endosomes in an EEA1-independent fashion. Together, these findings identify VP3 as the central orchestrator of endosomal remodeling during birnaviruses infection and uncover a conserved reliance on PtdIns(3)P-dependent endosomal platforms that are adapted through distinct host-specific interactions across two evolutionarily distinct birnaviruses.

## Results

### VP3-Es are hybrid compartments that feature early and late endosomal maturation markers

The persistent association of IBDV VP3 with PtdIns(3)P-positive compartments, both during ectopic expression and throughout infection (12), suggested that VP3-Es originate from the early endosomal compartment. However, VP3-Es differed markedly from canonical early endosomes. Whereas early endosomes are short-lived peripheral organelles that rapidly mature into late endosomes (14), VP3-Es were consistently localized in the perinuclear region and enriched in PtdIns(3)P (12), a lipid normally lost during endosomal maturation (15). These observations suggested that VP3 not only associates with early endosomes but also remodels them into distinct long-lived compartments that may contribute to the formation of IBDV replication compartments. To test this possibility, we characterized the identity of VP3-Es in avian QM7 cells stably expressing IBDV VP3 (QM7-VP3). For this purpose, cells were transiently transfected with fluorescently tagged markers of early, recycling, late, and terminal endosomal compartments and analyzed for their co-occurrence with VP3 puncta. As shown in Figure 1A, VP3-Es displayed a predominantly perinuclear distribution and were positive for the early endosomal markers Rab5 and PtdIns(3)P, consistent with our previous observations (12). However, VP3-Es failed to recruit Rab4, the small GTPase responsible for the formation of recycling vesicles from early endosomes. In addition, they lacked both Rab7 and Rab9, markers of the degradative and late retrograde trafficking routes, respectively (16). These findings indicate that VP3-Es retain selected features of early endosomes while remaining excluded from canonical recycling and maturation pathways. Furthermore, VP3-Es showed no association with Arl8, a small GTPase involved in the maturation of late endosomes into terminal endocytic compartments (17). In contrast, these compartments were positive for Lamp1 and Lamp3 (CD63) (Figure 1A, and Figure S1A), which are typically enriched in late and terminal endocytic compartments (18). Furthermore, the vast majority of VP3-Es showed Rab5 and PtdIns(3)P with Lamp1, but not Rab7, co-existing within the same compartments (Figure 1B). Together, these findings indicate that VP3-Es feature a hybrid endosomal identity, combining hallmarks of early and late endosomes while lacking key markers associated with progression toward the terminal degradative pathway (lack of Rab7 and Arl8b).

**Figure 1.**
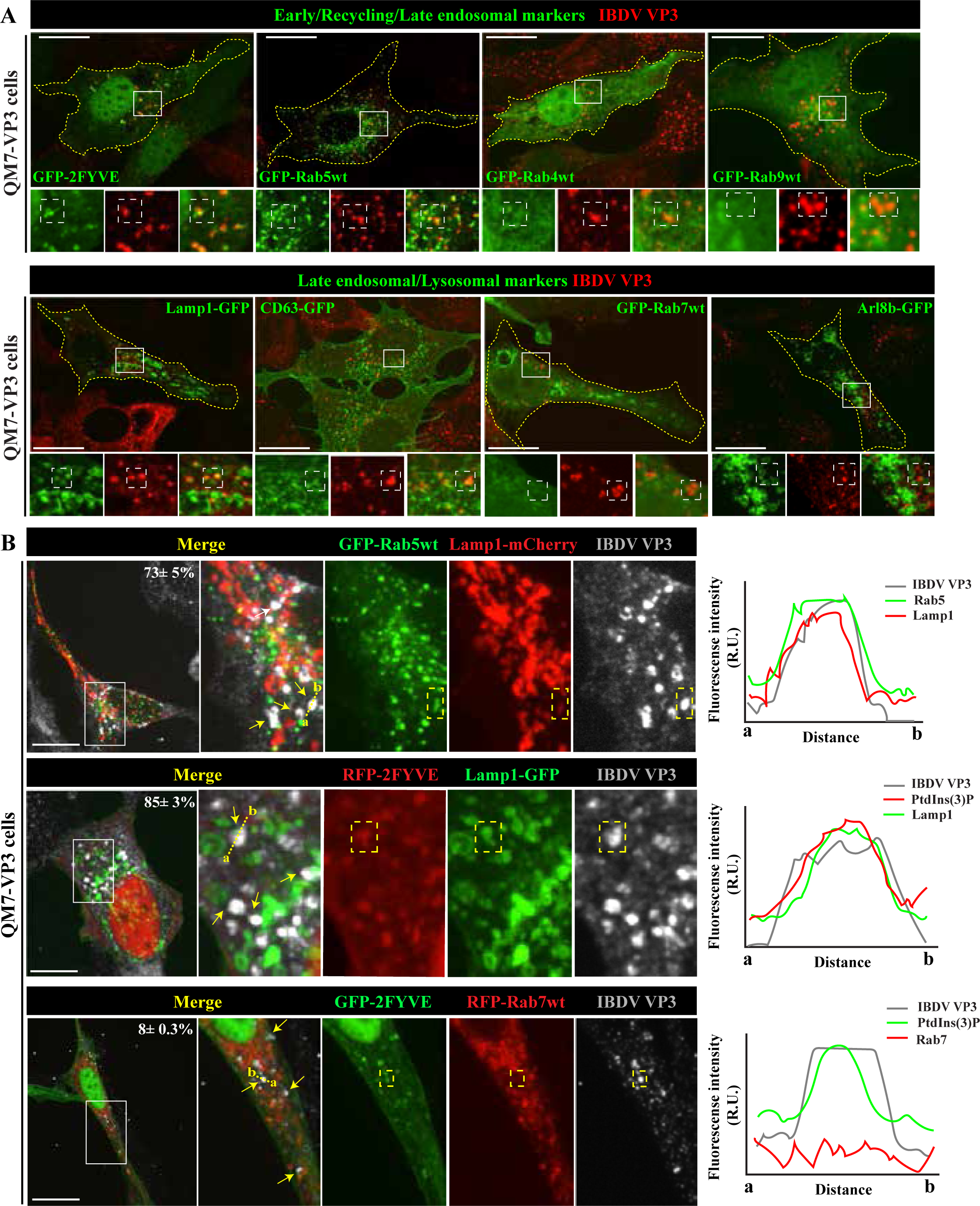
VP3-Es are hybrid endocytic compartments. (A) Spinning-disk confocal microscopy (SDCM) images of QM7-VP3 cells showing VP3 (red) and the indicated endocytic markers (green): GFP-2FYVE, GFP-Rab5wt, GFP-Rab4wt, GFP-Rab9wt, Lamp1-GFP, CD63-GFP, GFP-Rab7wt, and Arl8b-GFP. (B) SDCM images of QM7-VP3 cells showing VP3 (gray) together with combinations of early and late endocytic markers: GFP-Rab5wt/Lamp1-mCherry, RFP-2FYVE/Lamp1-GFP, and GFP-2FYVE/RFP-Rab7wt. Yellow arrows indicate VP3-Es positive for both early and late endocytic markers. Fluorescence intensity profiles along the dashed lines (a-b) are shown on the right. Percentages of VP3/early endocytic marker/late endocytic marker-positive compartments are shown as mean ± SD from three independent experiments. Magnified views show split-channel and merged views of representative VP3-Es. Dashed boxes indicate individual VP3-Es. Images are representative of three independent experiments. Scale bars, 10 μm.

Additionally, VP3-Es failed to acidify despite exhibiting late endosomal characteristics (Figures 2A and B and Figure S1 B and C), suggesting that VP3 disrupts endosomal acidification. We therefore examined the vacuolar ATPase (V-ATPase), whose activity depends on the reversible assembly of the cytosolic V1 domain with the membrane-embedded V0 domain (19). Using a fluorescent SidK-derived probe that specifically binds the V1 A subunit (20), we detected robust V1 recruitment to VP3-Es (Figure 2C). Thus, defective V1 recruitment is unlikely to account for the impaired acidification of these compartments. Next, we asked whether the defective acidification of VP3-Es resulted from loss of membrane integrity. To test this, we used the membrane damage reporters YFP-Galectin-3 (21) and GFP-Lysenin (W20A) (22). Neither probe localized to VP3-Es (Figures 2D and 2E), whereas both were robustly recruited to vesicles following permeabilization with L-leucyl-L-leucine methyl ester (LLOMe) (23) (Figure S1D). These findings indicate that impaired acidification of VP3-Es occurs in the absence of detectable endosomal membrane damage. VP3-Es also failed to undergo the Rab5-to-Rab7 conversion that defines canonical endosomal maturation. Consistent with this phenotype, VP3-Es lacked the MON1-CCZ1 complex (Figure 2F), which promotes Rab7 recruitment during endosome maturation (24). In contrast, VP3-Es were enriched in Rubicon (Figure 2G), a negative regulator of endosomal maturation that suppresses Rab7 activation (25). In line with these results, VP3-Es were resistant to chemical breakage by LLOMe, a disruptor molecule whose function depends on acidity and hydrolase activity (23) (Figure 2H). Collectively, these findings identify VP3-Es as hybrid, non-acidic endocytic compartments that retain features of both early and late endosomes while failing to acquire degradative properties.

**Figure 2.**
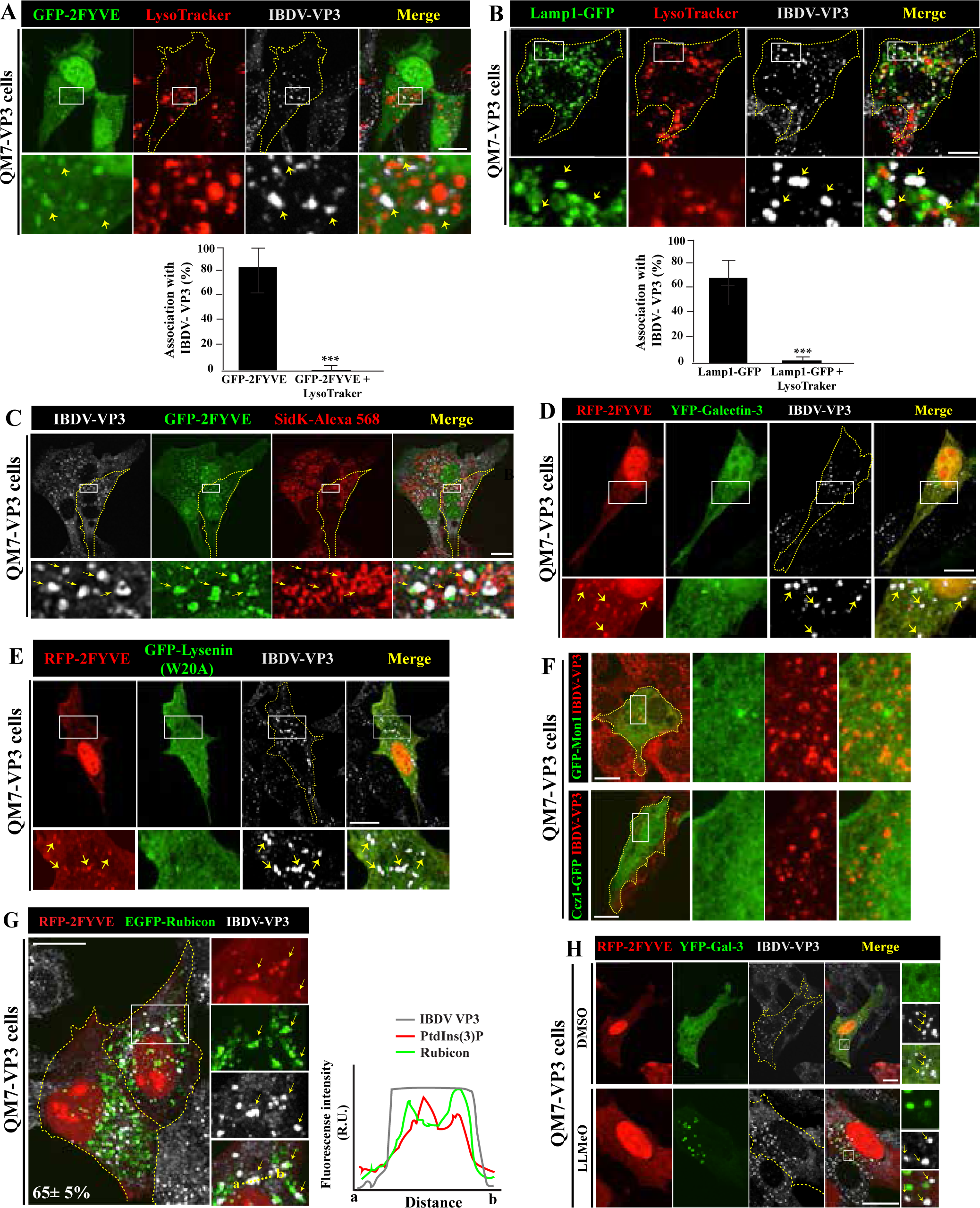
VP3-Es are non-acidic, intact compartments with halted maturation. (A,B) SDCM images of QM7-VP3 cells showing VP3 (gray), GFP-2FYVE or Lamp1-GFP (green), and LysoTracker (red). Bar graphs show the percentage of VP3-Es positive for the indicated endocytic markers and LysoTracker. (C) QM7-VP3 cells expressing GFP-2FYVE (green) and labeled with the V-ATPase probe SidK-Alexa 568 (red). (D,E) QM7-VP3 cells co-expressing RFP-2FYVE (red) and the membrane damage reporters YFP-galectin-3 or GFP-lysenin W20A (green). (F) QM7-VP3 cells expressing GFP-Mon1 or Ccz1-GFP (green). (G) QM7-VP3 cells co-expressing RFP-2FYVE (red) and EGFP-Rubicon (green). Yellow arrows indicate VP3-Es positive for both PtdIns(3)P and Rubicon. Percentages of VP3/PtdIns(3)P/Rubicon-positive compartments are shown as mean ± SD from three independent experiments. Fluorescence intensity profiles along the dashed lines (a-b) are shown on the right. (H) QM7-VP3 cells co-expressing RFP-2FYVE (red) and YFP-galectin-3 (green) following LLOMe-induced membrane damage. Yellow arrows point to VP3-Es resistant to chemical breakage. Magnified views show representative VP3-Es. Yellow arrows indicate representative VP3-Es where shown. Images are representative of three independent experiments. Scale bars, 10 μm.

### VP3 mediates PtdIns(3)P persistence on early endosomes

Time-lapse video microscopy of cells stably expressing fluorescently tagged VP3 ( acGFP-IBDV-VP3) revealed the emergence of VP3-Es at actively ruffling cell edges (Supplementary Video 1 and Figure 3A), consistent with recruitment to newly formed PtdIns(3)P-positive early endosomes. These compartments subsequently trafficked to and accumulated within the perinuclear region (Figure 3B and Figure S1E), recapitulating the distribution and PtdIns(3)P association previously observed for untagged IBDV VP3. Together, these observations suggested that VP3 captures nascent early endosomes and stabilizes PtdIns(3)P on their membranes, diverting them from the canonical maturation pathway. To test this hypothesis, we monitored PtdIns(3)P turnover on phagosomes, a tractable model of endosomal maturation. QM7 and QM7-VP3 cells expressing FcγR efficiently internalized IgG-opsonized latex beads (Figure 3C). Although newly formed phagosomes in both cell types acquired PtdIns(3)P, the loss of this lipid during phagosome maturation, monitored using GFP-2xFYVE, was markedly delayed in QM7-VP3 cells (Figure 3D).

**Figure 3.**
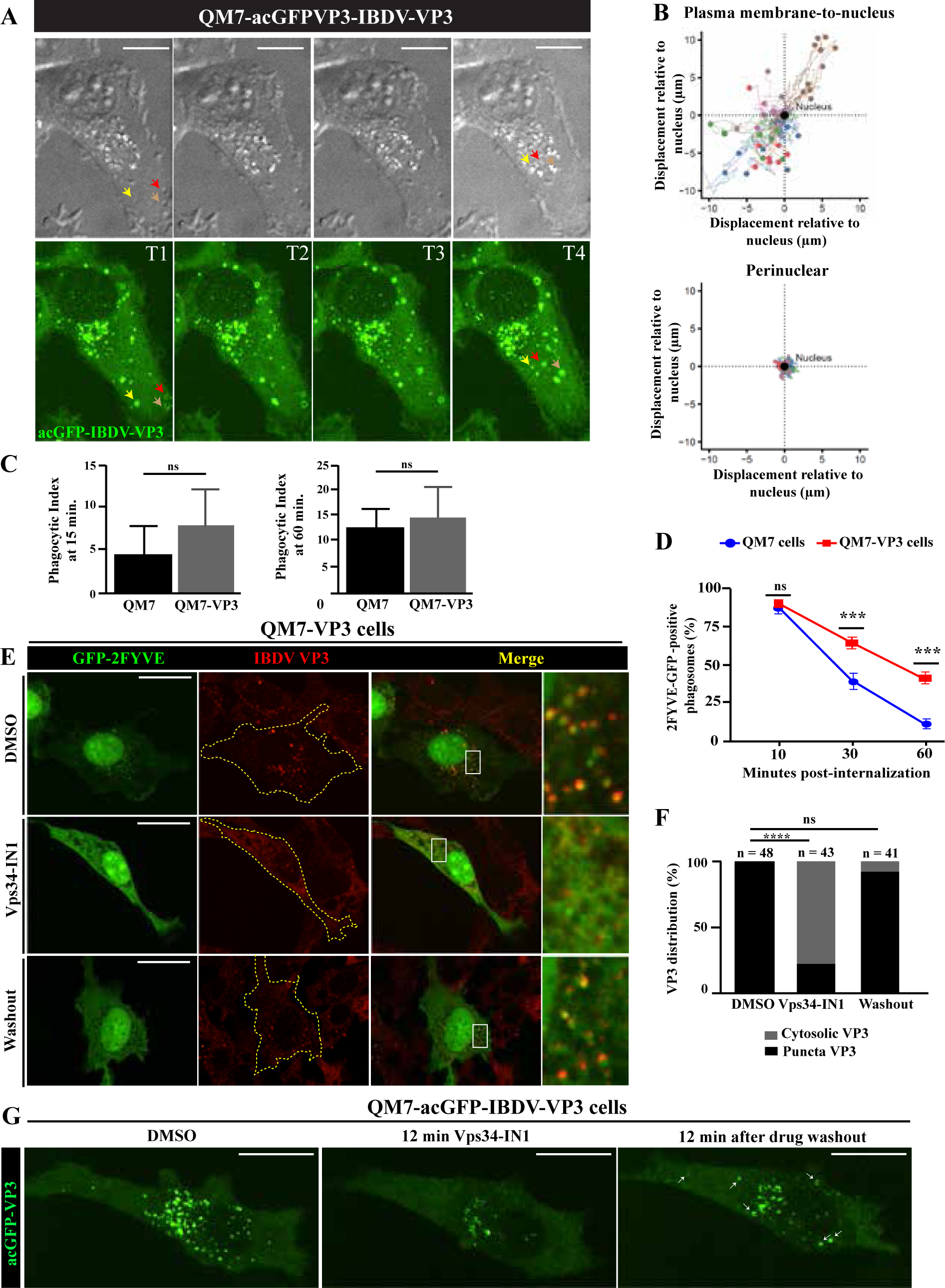
VP3 mediates PtdIns(3)P persistence on early endosomes. (A) Representative frames from time-lapse video microscopy of QM7-acGFP-IBDV-VP3 cells. T1, 1 min 15 s; T2, 1 min 26 s; T3, 1 min 58 s; T4, 3 min 46 s. Colored arrows indicate emerging VP3-Es at T1 and their subsequent displacement through T4. (B) Plots represent the trajectory analysis of individual VP3-Es tracked throughout the entire imaging period (25 min). (C) Phagocytic index of QM7 and QM7-VP3 cells transiently expressing myc-FcγRIIA. A synchronized phagocytosis assay was performed as described in Materials and Methods. Values represent mean ± SD from three independent experiments (ns, not significant). (D) Kinetics of PtdIns(3)P turnover in QM7 and QM7-VP3 cells co-expressing GFP-2FYVE and myc-FcγRIIA. Cells were fixed 10, 30, or 60 min after latex bead internalization and analyzed by SDCM. Graph shows the percentage of bead-containing phagosomes positive for PtdIns(3)P. Values represent mean ± SD from three independent experiments (ns, not significant; ***P < 0.001). (E,F) SDCM images and assessment of VP3 distribution in QM7-VP3 cells expressing GFP-2FYVE and treated with Vps34-IN1. Values represent SD from three independent experiments (ns, not significant; ****P < 0.0001). (G) Representative frames from time-lapse video microscopy of QM7-acGFP-IBDV-VP3 cells following Vps34-IN1 treatment and drug washout. White arrows indicate newly formed VP3-Es merging after drug removal. Images are representative of three independent experiments. Scale bars, 10 μm.

Because conversion of phosphatidylinositol to PtdIns(3)P by the class III PI3-kinase Vps34 is the primary source of endosomal PtdIns(3)P (18), we next examined whether maintenance of VP3-Es depended on continuous PtdIns(3)P synthesis. Treatment of QM7 cells co-expressing IBDV VP3 and GFP-2xFYVE with the reversible Vps34 inhibitor Vps34-IN1 caused the dissociation of both proteins from endosomal membranes and their redistribution to the cytosol (Figure 3 E, F). Removal of the inhibitor restored GFP-2xFYVE- and VP3-Es (Supplementary Video 2, Figure S2A and Figure 3G). The rapid loss of PtdIns(3)P following Vps34 inhibition indicates that the enzymatic pathways responsible for PtdIns(3)P turnover (26) remain active within the VP3-Es. These findings suggest that persistence of PtdIns(3)P on VP3-Es is maintained by sustained Vps34-dependent synthesis that counterbalances ongoing turnover. Collectively, these data indicate that VP3 captures early endosomes and remodels them into hybrid maturation-arrested organelles in which PtdIns(3)P persistence depends on continuous lipid production.

### The PtdIns(3)P effector ESCPE-1 promotes retrograde trafficking of VP3-Es and is essential for IBDV replication

The directional trafficking and perinuclear accumulation of VP3-Es in QM7 cells recapitulated the juxtanuclear positioning of RNP-bearing endosomes observed during IBDV infection, suggesting that VP3 engages the retrograde trafficking that contribute to the formation of birnaviruses replication compartments (8, 9, 12, 13). Consistent with this hypothesis, brief treatment of QM7-acGFP-VP3 cells with dynarrestin, an inhibitor of the microtubule motor dynein (27), caused VP3-Es to lose their perinuclear localization and disperse throughout the cytoplasm (Figure 4A), indicating that dynein-dependent retrograde transport is required for their positioning. We therefore focused on ESCPE-1, a PtdIns(3)P-binding sorting complex composed of SNX1/2 and SNX5/6 heterodimers that drives endosome-to-Golgi retrograde trafficking (28). Owing to technical limitations in avian cells, these experiments were conducted in HeLa cells after verifying that VP3-Es displayed the same trafficking behavior observed in QM7 cells. Specifically, VP3-Es formed at the cell periphery and rapidly trafficked toward the juxtanuclear region, where their motility became markedly reduced (Supplementary Video 3 and Figure S2B). As shown in Figure 4B and Figure S2C, the SNX proteins, which displayed a predominantly perinuclear distribution in control HeLa cells, were recruited to VP3-Es in acGFP-VP3-expressing cells. Importantly, the depletion of PtdIns(3)P redistributed both SNX1 and VP3 to the cytoplasm (Figure 4C), indicating that PtdIns(3)P is required for ESCPE-1 recruitment to VP3-Es. To assess the role of ESCPE-1 in VP3-Es trafficking, we disrupted the complex in SNX5/6-knockout (SNX5/6 KO) HeLa cells by silencing SNX1 and SNX2 (Figure S2 D, E). Loss of ESCPE-1 function was verified using the cation-independent mannose 6-phosphate receptor (M6PR), a known ESCPE-1 cargo (29) (Figure 4D, upper panel). Similarly, the SNXs depletion disrupted the characteristic perinuclear distribution of VP3-Es, resulting in their redistribution throughout the cytoplasm (Figure 4D, lower panel). Together, these results indicate that sustained PtdIns(3)P on VP3-Es promotes ESCPE-1 recruitment, thereby driving their retrograde transport and perinuclear accumulation. We then tested whether ESCPE-1 is required for IBDV replication. To minimize effects on cell viability, we first depleted SNX1 by siRNA. SNX1 depletion markedly reduced VP3 accumulation in infected cells relative to controls (Figures 4E and F). Given the functional redundancy between SNX1 and SNX2(29), both proteins were subsequently depleted simultaneously. Efficient SNX1/SNX2 silencing nearly abolished VP3 expression and dramatically reduced viral progeny production (Figure 4G and H). Together, these findings demonstrate that the ESCPE-1 complex is recruited to and mediates the retrograde trafficking of VP3-Es and acts as a critical host factor for IBDV replication.

**Figure 4.**
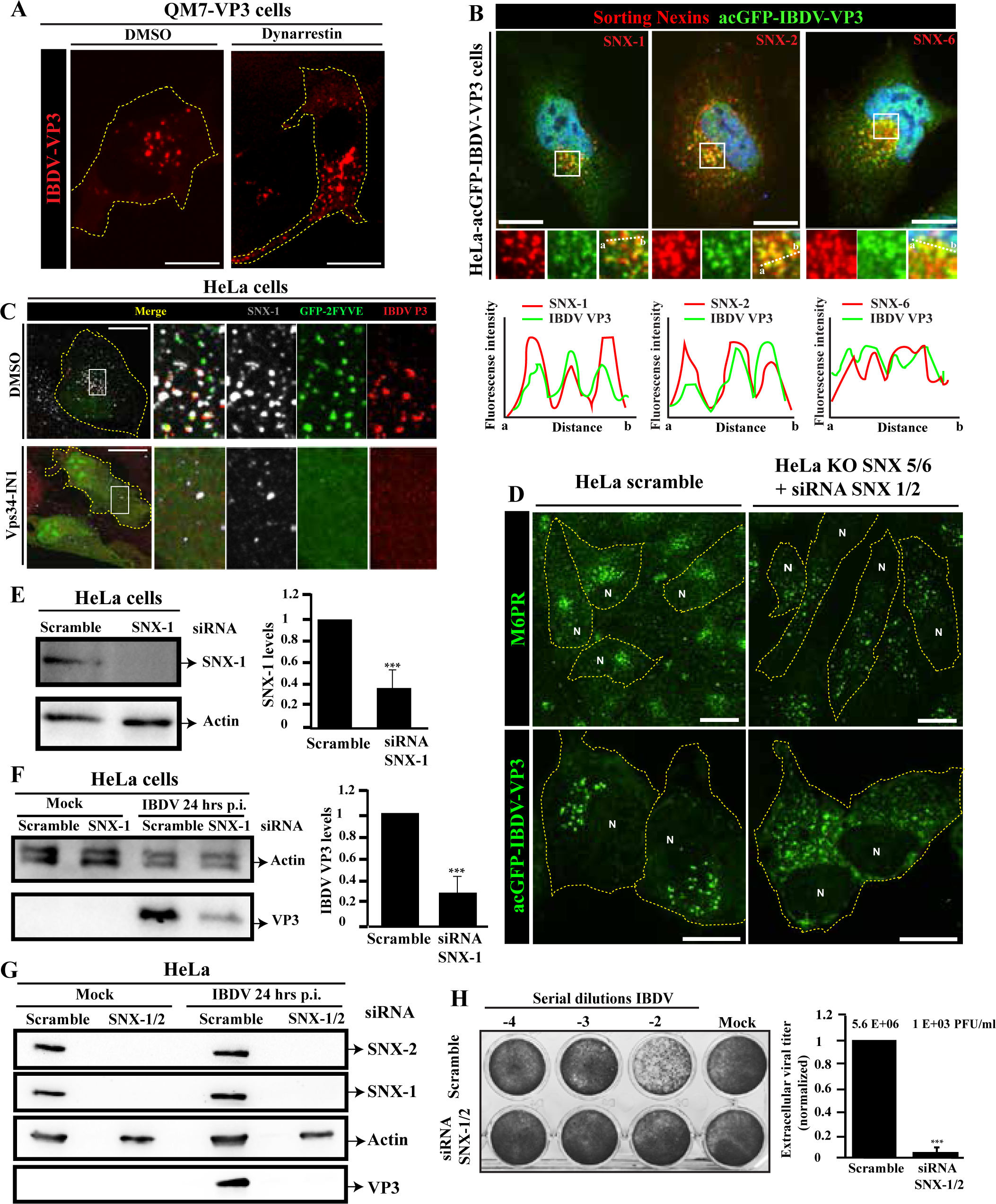
ESCPE-1 promotes retrograde trafficking of VP3-Es and is essential for IBDV replication. (A) SDCM images of QM7-VP3 (red) cells treated with Dynarrestin. (B) SDCM images of acGFP-IBDV-VP3 HeLa cells (green) stained for SNX1, SNX2, or SNX6 (red). Magnified views show representative VP3-Es. Fluorescence intensity profiles along dashed lines (a-b) are shown below each panel. (C) SDCM images of HeLa-VP3 (red) cells expressing GFP-2FYVE and treated with Vps34-IN1, followed by immunostaining for SNX1(gray) and VP3 (red). Magnified views show representative VP3-Es. (D) SDCM images of control or SNX1/2/5/6-depleted HeLa cells stained for M6PR (green) or expressing acGFP-IBDV-VP3 (green). N, nucleus. (E,F) Immunoblot analysis of SNX1-depleted HeLa cells in the absence (E) or presence (F) of IBDV infection. Quantification is shown as mean ± SD from three independent experiments (**P < 0.01). (G) Immunoblot analysis of SNX1/SNX2-depleted HeLa cells following IBDV infection. (H) Plaque assay and quantification of extracellular viruses from SNX1/SNX2-depleted HeLa cells. Normalized viral titers are shown as mean ± SD from three independent experiments (***P < 0.01); raw titers are indicated above the bars. Images and immunoblots are representative of three independent experiments. Scale bars, 10 μm.

### VP3-Es tether to the Golgi complex

Given that VP3-Es recapitulate the trafficking behavior of RNP-hijacked endosomes (9, 13), we sought to determine whether VP3 is the key factor driving the association of these endosomes with the Golgi complex.

On-section correlative light and electron microscopy (CLEM) revealed VP3-Es of heterogeneous size positioned adjacent to the trans-Golgi network, with several vesicles closely apposed to the Golgi cisternae (Figure 5A). Consistent with this observation, disruption of Golgi function with Golgicide A, an inhibitor of the ARF-GEF GBF1 (30), dispersed VP3-Es from their characteristic perinuclear location. Likewise, Brefeldin A, which induces Golgi collapse into the endoplasmic reticulum and promotes fusion of the trans-Golgi network with early endosomes (31), markedly reduced the abundance of VP3-Es (Figure 5B). Together, these findings indicate that the perinuclear accumulation of VP3-Es depends on an intact Golgi apparatus. Using the proximal and distal Golgi markers SialT2-YFP and GalNAcT-YFP, respectively (32), we found VP3-Es closely associated with both proximal and distal Golgi cisternae, with a stronger association observed at the distal Golgi (Figure 5C). Importantly, this interaction persisted after nocodazole treatment, which disrupts microtubule polymerization and fragments the Golgi into dispersed ministacks (33) (Figure 5D). Under these conditions, VP3-Es remained associated with Golgi ministacks, consistent with previous observations made during viral infection (13) and underscoring VP3’s intrinsic ability to drive Golgi tethering. The maintenance of this association phenotype despite microtubule disruption indicates that VP3-mediated Golgi association is microtubule-independent and relies on a stable interaction of VP3-Es with Golgi membranes.

**Figure 5.**
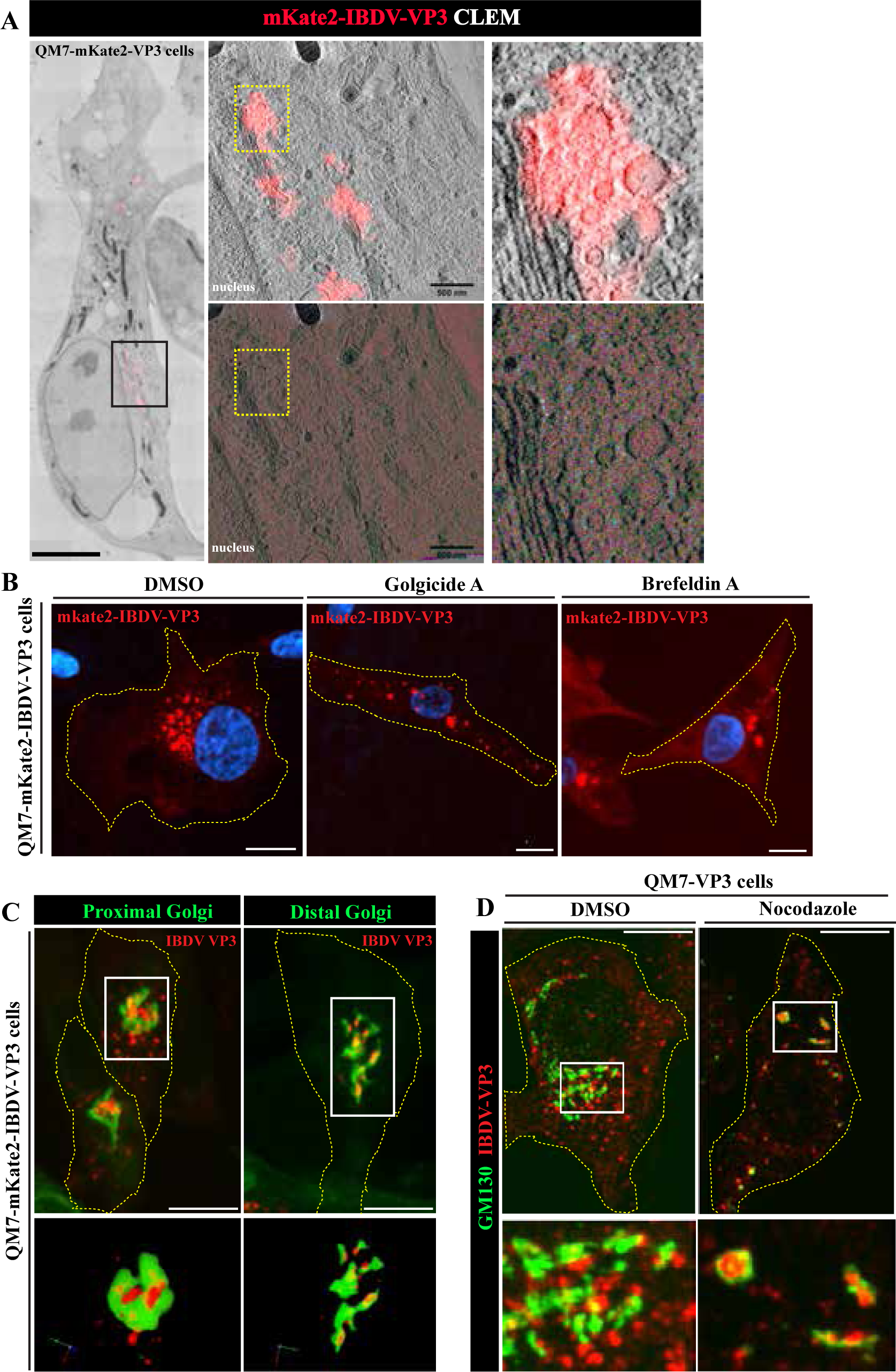
VP3-Es tether to the Golgi complex. (A) Correlative light and electron microscopy of QM7-mKate2-IBDV-VP3 cells. The boxed region identifies the area of interest; overlays of fluorescence and tomographic sections are shown at right. Magnified views highlight mKate2-VP3 signal associated with single-membrane vesicles adjacent to the trans-Golgi network. (B) SDCM images of QM7-mKate2-IBDV-VP3 (VP3, red) cells treated with DMSO, Golgicide A, or brefeldin A. (C) SDCM images of QM7-mKate2-IBDV-VP3 cells (red) expressing the proximal Golgi marker SialT2-YFP or the distal Golgi marker GalNAcT-YFP (green). Boxed regions were used for 3D reconstructions. (D) SDCM images of QM7-VP3 cells treated with DMSO or nocodazole. Magnified views show VP3-Es-Golgi contact areas. Images are representative of three independent experiments unless otherwise indicated. Panel A is representative of six tomograms. Scale bars, 10 μm.

### The VP3-PtdIns(3)P-endosome triad is central to birnaviruses replication

The findings above identify VP3 as the orchestrator of endosomal remodeling during IBDV replication. We therefore asked whether the VP3-dependent exploitation of PtdIns(3)P-positive endosomes is conserved in other birnaviruses. To address this question, we extended our analysis to the IPNV (34). Because the replication cycle of IPNV remains virtually unknown, we first examined the spatiotemporal dynamics of IPNV RNPs in infected CHSE/F fish cells using IPNV VP3 and dsRNA immunostaining for confocal imaging. As shown in Figure 6A, VP3 puncta were readily detected in the cytoplasm as early as 4 hpi. By 8 hpi, viral dsRNA became detectable and both VP3 and dsRNA accumulated predominantly in the perinuclear region (Figures 6A and 6B, middle panels). At later stages of infection (24 hrs p.i.), VP3 and dsRNA were redistributed into large cytoplasmic aggregates that extended beyond the perinuclear region (Figures 6A and B, right panels). Co-immunostaining of these RNP components revealed VP3/dsRNA positive puncta at the perinuclear area of infected cells (Figure 6C). Importantly, both RNP components localized to PtdIns(3)P-positive vesicles (Figure 6D), recapitulating the organization previously observed for IBDV. These findings suggested that PtdIns(3)P-positive endosomes also support IPNV replication. To test this possibility, we depleted endosomal PtdIns(3)P by treating infected CHSE/F cells with Vps34-IN1 (Figure S3A and B) and assessed its impact on viral replication. As shown in Figures 6E and 6F, depletion of PtdIns(3)P profoundly impaired IPNV replication, reducing VP2 and VP3 accumulation as well as intracellular and extracellular viral titers. Furthermore, viral dsRNA levels and transcript abundance were likewise significantly decreased in Vps34-IN1-treated cells (Figure S3C-D). Collectively, these data identify PtdIns(3)P-positive endosomes as essential replication platforms for IPNV and reveal a conserved VP3-PtdIns(3)P-endosome triad underlying birnaviruses replication.

**Figure 6.**
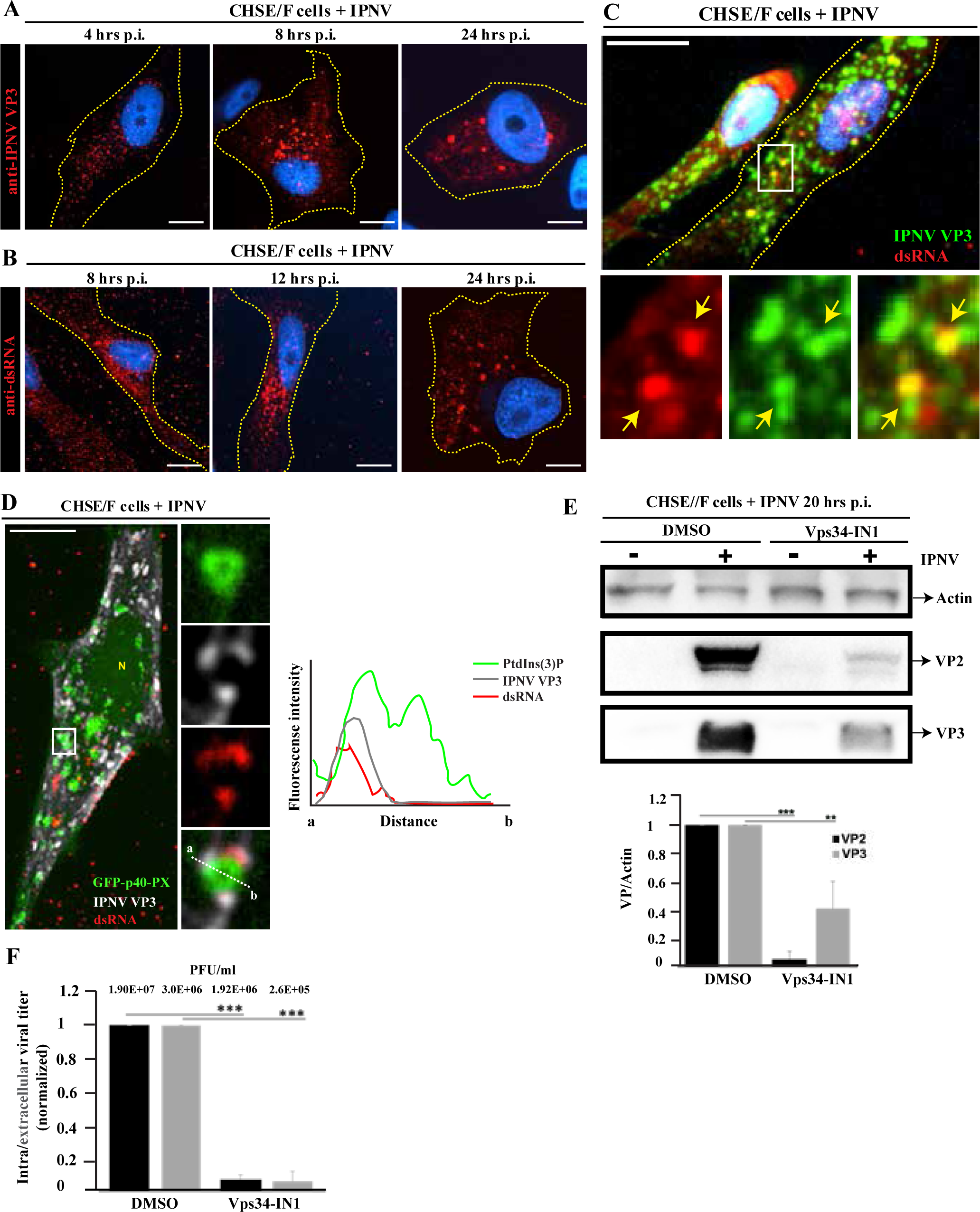
VP3-PtdIns(3)P-endosome triad is central to birnaviruses replication. (A,B) SDCM time-course analysis of CHSE/F cells infected with IPNV (MOI=5 PFU/cell), showing VP3 expression at 4, 8, and 24 h postinfection (hpi) (A) or dsRNA accumulation at 8, 12, and 24 hpi (B). (C) SDCM images of IPNV-infected CHSE/F cells at 12 hrs p.i. immunostained for VP3 (green) and dsRNA (red). Magnified views show representative VP3/dsRNA-positive puncta at the perinuclear region. Nuclei are shown in blue. (D) SDCM images of CHSE/F cells expressing the PtdIns(3)P biosensor GFP-p40-PX and infected with IPNV. Cells were stained for VP3 and dsRNA at 12 hpi. Magnified views show representative PtdIns(3)P/VP3/dsRNA-positive puncta. Fluorescence intensity profiles along the dashed lines (a-b) are shown on the right. N, nucleus. (E) Immunoblot analysis of IPNV-infected CHSE/F cells treated with Vps34-IN1. Quantification is shown as mean ± SD from three independent experiments (**P < 0.05; ***P < 0.01). (F) Intra- and extracellular virus titers from IPNV-infected CHSE/F cells treated as in (E). Values represent mean ± SD from three independent experiments (***P < 0.01); raw titers are indicated above the bars. Images and immunoblots are representative of three independent experiments. Scale bars, 10 μm.

### EEA1 mediates targeting of IBDV VP3 to early endosomes

The findings above support a conserved replication strategy for birnaviruses based on VP3-dependent hijacking of PtdIns(3)P-positive endosomes. Consistent with this model, structural predictions identified conserved electropositive regions in IPNV VP3 homologous to the endosome-targeting PATCH2 domain of IBDV VP3 (Figure S4A), suggesting a common mechanism of endosomal targeting. To determine whether this process involves direct PtdIns(3)P recognition, we examined the binding of recombinant IBDV VP3 and IPNV VP3 to phosphoinositides. Consistent with previous reports (8, 12), IBDV VP3 displayed broad, low-specificity phosphoinositide binding, whereas IPNV VP3 showed only background-level binding to PtdIns(3)P (Figure 7A, upper immunoblots). Likewise, semiquantitative lipid-overlay assays using serial PtdIns(3)P dilutions failed to detect specific bindings by either protein (Figure 7A, lower immunoblots). These findings were confirmed using PtdIns(3)P-PIPosomes. As shown in Figure 7B, both VP3 proteins associated equally with control and PtdIns(3)P-PIPosomes. Together, these results argue against direct PtdIns(3)P recognition by VP3 and instead suggest recruitment through a PtdIns(3)P-binding host factor. VP3 immobilized on magnetic beads was therefore used as bait in pull-down assays from post-nuclear cell lysates, followed by mass spectrometry analysis (LC-ESI-MS/MS) to identify interactors (Figure S4B). Because proteomic annotation of avian and fish cells remains limited, we performed these analyses in HeLa cells using purified IBDV VP3 as bait. Among the proteins identified by mass spectrometry, the early endosomal PtdIns(3)P effector EEA1 emerged as the top candidate interactor (Figure S4C). EEA1 is a Rab5- and PtdIns(3)P-binding tethering factor that is specifically recruited to early endosomal membranes (35). These properties make EEA1 a strong candidate for mediating VP3 recruitment to PtdIns(3)P-positive endosomes. To validate the proteomic findings, we performed pull-down assays followed by immunoblotting. As shown in Figure 7C and Figure S4D, IBDV VP3 robustly associated with ectopically expressed tagged-EEA1, whereas no interaction was detected for IPNV VP3. Consistent with these results, EEA1 extensively colocalized with IBDV VP3-positive vesicles, whereas IPNV VP3 showed no detectable association with EEA1-positive endosomes and remained largely cytoplasmic (Figure 7D). Together, these findings indicate that IBDV VP3, but not IPNV VP3, targets PtdIns(3)P-positive endosomes through interaction with EEA1. This result raised the possibility that the interaction between VP3 and EEA1 may be host dependent. To test this, we performed pull-down assays using IBDV VP3 and IPNV VP3 as bait and post-nuclear lysates from fish (CHSE/F and ZKS) and avian (QM7) cells. As shown in Figures 7E-F and Figure S4E, IBDV VP3 efficiently captured endogenous EEA1 from both fish and avian cells, whereas IPNV VP3 failed to detectably interact with EEA1 in either cell line. Consistent with these biochemical data, IBDV VP3 formed perinuclear punctate structures in both fish and avian cells, whereas IPNV VP3 formed puncta only in fish cells and remained largely diffused in the cytoplasm of avian cells (Figure 7G). Together, these findings reveal distinct endosomal targeting mechanisms employed by VP3 orthologues. In IBDV VP3, association to PtdIns(3)P-positive endosomes is mediated by EEA1, a process conserved across human, avian, and fish cells. In contrast, another PtdIns(3) effector aside of EEA1 is required for the IPNV VP3 to associate to PtdIns(3)P-positive endosomes. This effector is yet to be identified.

**Figure 7.**
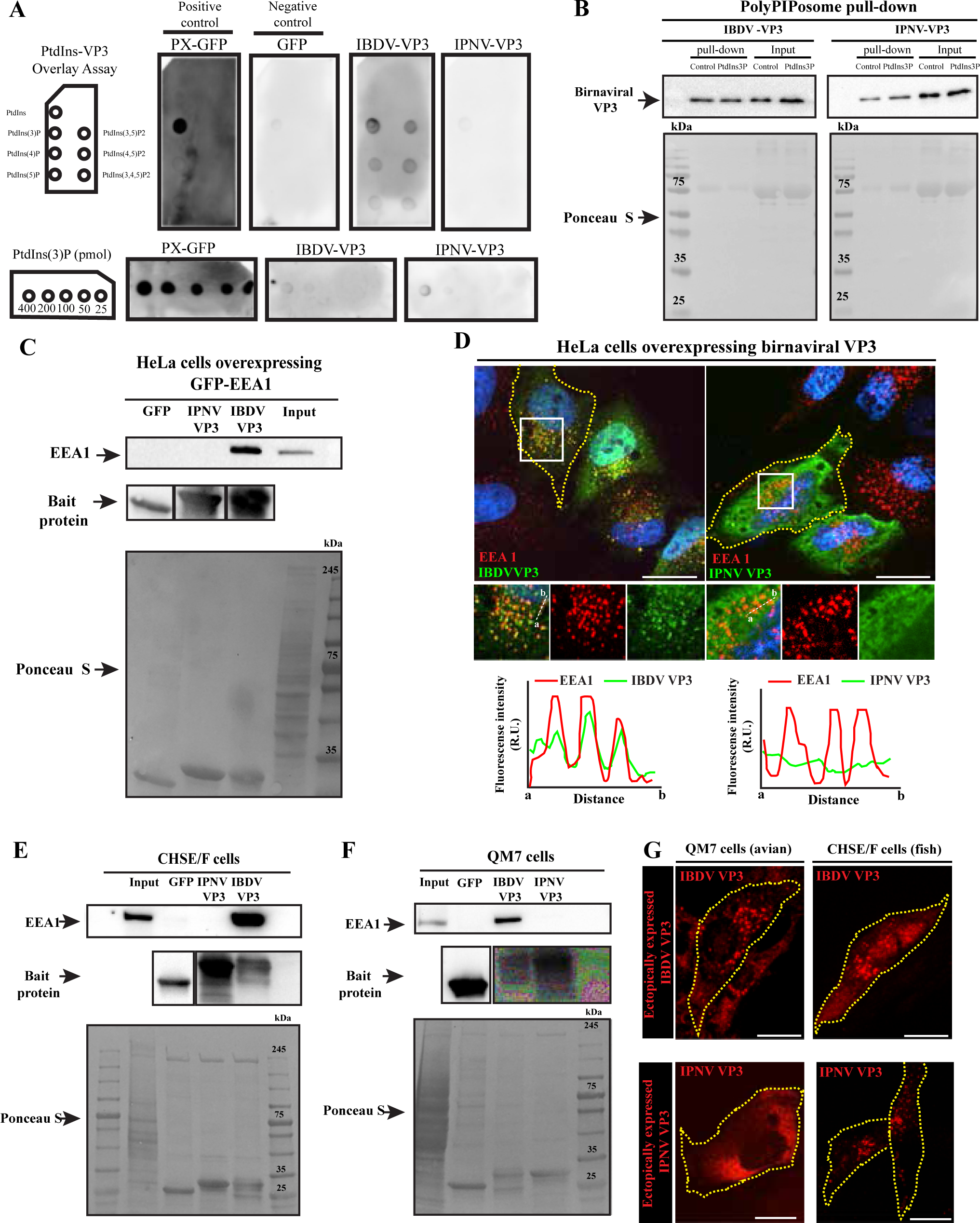
EEA1 mediates targeting IBDV VP3 to early endosomes. (A) Lipid-protein overlay assays using the indicated phosphoinositide species or increasing concentrations of PtdIns(3)P. Purified PX-GFP (positive control), GFP (negative control), IBDV-VP3, and IPNV-VP3 proteins were incubated with lipid strips as described in Materials and Methods. (B) Pull-down assays using control or PtdIns(3)P-containing PIPosome lipid nanoparticles and purified IBDV-VP3 or IPNV-VP3 proteins. (C) Pull-down assays performed with protein-coated magnetic beads and lysates from HeLa cells expressing GFP-EEA1. GFP served as a heterologous control. (D) SDCM images of HeLa cells expressing IBDV-VP3 or IPNV-VP3 and stained for EEA1. Magnified views show representative VP3/EEA1-positive compartments. Nuclei are shown in blue. Fluorescence intensity profiles along the dashed lines (a-b) are shown below each panel. (E,F) Pull-down assays performed with protein-coated magnetic beads and lysates from CHSE/F or QM7 cells. GFP served as a heterologous control. (G) SDCM images of QM7 and CHSE/F cells expressing IBDV-VP3 or IPNV-VP3. Cells were processed for immunofluorescence detection of VP3 proteins. Immunoblots and images are representative of three independent experiments. Scale bars, 10 μm.

## Discussion

ROs are typically established by viral proteins that target and remodel specific host membranes, a process extensively studied in +ssRNA viruses (11). While many of these viruses rely on transmembrane or hydrophobic membrane-binding domains for this purpose, others exploit phosphoinositides to anchor replication complexes and remodel organelle identity (36–41). VP3 differs fundamentally from these paradigms. Despite lacking recognizable membrane-binding motifs, VP3 associates robustly with PtdIns(3)P-positive endosomes, a process previously attributed to direct lipid recognition through its PATCH2 domain (42). Using VP3 expressed in a prokaryotic system and complementary biochemical and cellular approaches, we found no evidence that either IBDV or IPNV VP3 directly binds PtdIns(3)P. Instead, our findings identify the PtdIns(3)P effector EEA1 as the host factor that recruits IBDV VP3 to early endosomal membranes, thereby reconciling the dependence of VP3 on PtdIns(3)P-positive endosomes with the absence of detectable lipid binding. Our data suggest that VP3 interacts directly with EEA1 since no other known EEA1-interacting proteins, which could bridge this binding, were detected in the pulldown-MS. Notably, IPNV VP3 associated with endosomes independently of EEA1, indicating that VP3 orthologues have evolved distinct host-factor interactions to access a conserved PtdIns(3)P-positive replication platform.

Beyond endosomal targeting, VP3 drives the formation of long-lived, non-acidic hybrid endocytic organelles that accumulate in the perinuclear region and associate with the Golgi complex. These compartments retain features of both early (Rab5, PtdIns(3)P, EEA1) and late (Lamp proteins) endosomes but fail to acquire a degradative identity, suggesting that VP3 arrests endosomal maturation, at least in part, through impaired Rab5-to-Rab7 conversion. In this regard, the localized enrichment of Rubicon on VP3-Es suggests it may act as the host factor preventing Rab7 activation on the endosomal membrane (25).

A defining feature of VP3-Es is the persistence of PtdIns(3)P, which normally turns over rapidly during endosomal maturation (43). The rapid loss of PtdIns(3)P following Vps34 inhibition in QM7-VP3 cells indicates that the lipid turnover remains active and suggests that persistent PtdIns(3)P is maintained through sustained Vps34-dependent synthesis, potentially favored by the neutral luminal pH of VP3-Es (44). VP3 dual role of hijacking and reprograming host endosomes through the modulation of PtdIns(3)P dynamics parallels those described in a group of +ssRNA viruses that also targets phosphoinositides metabolism (45–47). Similarly, the tomato bushy stunt virus (TBSV) targets PtdIns(3)P for replication, but through a different mechanism than VP3. The TBSV p33 protein recruits the Vps34 causing PtdIns(3)P accumulation on peroxisomal and ER membranes to form ROs (48). Furthermore, the hepatitis C virus (HCV) protein NS5A and the enteroviral protein 3A promote the accumulation of PtdIns(4)P in the endoplasmic reticulum (ER) by recruiting the phosphatidylinositol 4-kinase IIIα (PI4KIIIα) and PI4KIIIβ, respectively (41, 49). These commonalities between IBDV and +ssRNA viruses are consistent with the hypothesis of birnaviruses as evolutionary intermediaries between distant RNA families.

Many viruses exploit endosome-to-Golgi trafficking pathways to support infection (50), and here we show that IBDV co-opts this route through VP3. By promoting PtdIns(3)P persistence, VP3 recruits the ESCPE-1 complex and drives dynein-dependent retrograde trafficking of hijacked endosomes towards the Golgi complex, where they accumulate and tether. This reprogramming of endosomal dynamics is essential for productive IBDV infection. Notably, while viral exploitation of the PtdIns(3)P-SNX axis has been reported in TBSV for ROs biogenesis, the strategy employed by VP3 appears mechanistically distinct. Whereas TBSV uses the non-structural protein p33 to stabilize pre-existing replication membranes (51), IBDV employs a structural component of the RNP to capture, remodel, and reposition host endosomes. Although the molecular basis of Golgi tethering remains unresolved, the enrichment of the trans-Golgi tethering proteins golgin-97 and golgin-245 in the VP3 interactome (data not shown), together with the apparent absence of VP3-Es-Golgi fusion, suggests that VP3 promotes stable endosome-Golgi contacts. Such contacts could provide a platform for the spatial and temporal coordination of viral replication and particle assembly (13).

By extending our analysis to the fish pathogen IPNV, we show that endosome-based replication is conserved across two representative members of the *Birnaviridae* family. In fish cells infected with IPNV, the replication machinery—specifically dsRNA and VP3— associates with PtdIns(3)P-enriched compartments localized within the perinuclear region. Furthermore, depletion of endosomal PtdIns(3)P severely compromised IPNV infectivity, thus identifying the VP3-PtdIns(3)P-endosome triad as a conserved feature of birnaviruses infection. Despite this conservation, the mechanisms of endosomal targeting have diverged. Whereas IBDV VP3 engages PtdIns(3)P-positive endosomes through EEA1 across multiple cell species, IPNV VP3 utilizes an EEA1-independent mechanism that appears restricted to piscine cells.

Collectively, our findings establish VP3 as a master organizer of birnaviruses replication and reveal how related birnaviruses have evolved distinct host-factor dependencies while preserving a common endosomal replication strategy. The discovery that avibirnavirus VP3 exploits EEA1 to associate with early endosomes is especially notable because, to our knowledge, no viral protein has previously been shown to directly co-opt this highly conserved endocytic tether. Future studies investigating the roles of EEA1, golgins, and SNARE proteins should provide further insight into how birnaviruses reprogram host endomembrane systems to establish replication-competent organelles.

## Materials and Methods

### Cell culture

QM7 (ATCC CRL-1962), HeLa (ATCC CCL-2), HeLa SNX5/6 knockout, 293^EL^, CHSE/F (formerly CHSE-214), and ZKS cells were maintained under standard conditions. QM7 and HeLa cells were cultured in DMEM supplemented with 10% FBS; QM7 medium additionally contained 25 mM HEPES. Stable VP3-expressing cell lines (untagged, mKate2-tagged, or acGFP-tagged VP3) were maintained in the presence of G418 (0.5 mg/mL for QM7 and 1 mg/mL for HeLa). Mammalian and avian cells were incubated at 37°C with 5% CO. CHSE/F cells were cultured in Leibovitz’s L-15 medium supplemented with 10% FBS at 20°C, or at 14°C during infection experiments. HeLa SNX5/6 knockout, 293EL, and ZKS cells were cultured as previously described (52). See supplementary information for more details on culture conditions and stable cell line development.

### Virus propagation and quantification

IBDV (Soroa strain) was propagated in QM7 cells and IPNV (Sp strain) in CHSE/F cells at an MOI of 0.05 PFU/cell. Following complete cytopathic effect, virus-containing lysates were subjected to three freeze-thaw cycles, clarified by centrifugation, and concentrated by PEG 8000 precipitation. IPNV preparations were further purified by sucrose cushion ultracentrifugation. IBDV titers were determined by plaque assay in QM7 cells, whereas IPNV titers were quantified by TCID endpoint dilution in CHSE/F cells and converted to PFU/mL using the Reed-Muench method. More details in SI Appendix.

### DNA constructs and transfection methods

Expression constructs encoding VP3, endosomal and lysosomal markers, phosphoinositide probes, and trafficking regulators were obtained from published sources, Addgene or generous gifts from collaborators, as detailed in supplementary information. Additional constructs generated for this study included fluorescently tagged IBDV VP3 and IPNV VP3 expression plasmids. HeLa and QM7 cells were transfected with Lipofectamine 3000, whereas CHSE/F cells were transfected with FuGENE HD and analyzed 24 h post-transfection. See SI Appendix for full details.

### Antibodies and other reagents

Commercial antibodies against endosomal and lysosomal markers, sorting nexins, Golgi proteins, viral proteins, dsRNA, GFP, and actin were obtained from established vendors or generated as custom polyclonal antisera against IBDV VP3, IPNV VP3, and purified IPNV virions. Fluorescently labeled and HRP-conjugated secondary antibodies were used for immunofluorescence and immunoblotting, respectively. Additional reagents included LysoTracker Red, fluorescent SidK, LLOMe, Brefeldin A, Golgicide A, Nocodazole, VPS34-IN1, and Dynarrestin. Full details of antibodies, reagents, and labeling procedures are provided in SI Appendix.

### siRNA-mediated knowckdown and IBDV infections

HeLa cells were transfected with siRNAs targeting **SNX1**, **SNX2**, or a non-targeting control using Lipofectamine RNAiMAX. Forty-eight hours post-transfection, cells were infected with IBDV at an MOI of 1 PFU/cell for 1 h. After virus adsorption, inocula were removed, cells were washed, and infection was allowed to proceed for 24 h. Samples were then harvested for immunoblot analysis and intracellular/extracellular virus titration.

### Treatment with pharmacological inhibitors

Vps34 activity was inhibited with VPS34-IN1 (1 µM) in QM7 and CHSE/F cells. Effects on VP3 localization, PtdIns(3)P production, and IPNV replication were assessed by fluorescence microscopy, immunoblotting, and viral titration. Golgi and microtubule function were perturbed in QM7 cells expressing mKate2-VP3 using Golgicide A, Brefeldin A, Nocodazole, or Dynarrestin, followed by fluorescence imaging.

### SDS-PAGE and western blotting for siRNA-mediated knock down, CRISP-Cas9 K0 and infection experiments

Cell monolayers were scrapped in complete media, centrifuged once at 300× g and washed twice with PBS 1X. Cells were lysed in NDP-40 lysis buffer supplemented with 1%Protease and phosphatase inhibitor cocktail (Ref: PPC1010-5M; Milipore-Sigma, Canada) for 10 minutes on ice, after which samples were centrifuged at 17,000g to obtain post-nuclear cell lysates. Cleared samples were mixed with 2x Laemmli sample buffer (Ref:1610737EDU; Bio-rad, Canada) and DL-Dithiothreitol solution (DTT) (Ref 43816; Milipore-Sigma, Canada) added to a final concentration of 2mM. The solution was heated at 95°C for 10 min and SDS-PAGE was performed on 10 or 12% polyacrylamide gels, and proteins transferred to nitrocellulose membranes using a semi-dry transfer system. membranes wash 3 times with distilled water and blocked with 10% skim milk in PBS 1X ON at 4°C. Membranes were washed twice with PBS 1X prior to incubation with the following antibodies: Purified mouse anti-sorting nexin 1 (Ref: 611482; 1/500), purified mouse monoclonal anti-sorting nexin 2 (Ref: 611308; 1/500), mouse monoclonal anti-sorting nexin 5 antibody (F-11) (Ref: sc-515215 ; 1/500), mouse monoclonal anti-sorting nexin 6 antibody (D-5) (Ref: sc-365965; 1/500), rabbit polyclonal anti-α-actin-1 antibody (Ref: A2066; 1/1000), customized rabbit polyclonal anti-IBDV VP3 (1/3000), anti-IPNV Vp2 protein mAb (clone 1B3/E10) (Ref: FM130CY-5; 1/1000), Mouse anti-IPNV VP3 protein, (clone 6E1/F2) (Catalog No. FM-130DZ-5; 1/1000), customized rabbit polyclonal anti-IPNV particles (1/5000), customized rabbit polyclonal anti-IPNV VP3 (1/2000). After three washings with PBS–0.05% Tween 20, for 15 min each, membranes were incubated with the corresponding HRP-conjugated secondary antibodies described above. Membranes were washed as described above and the corresponding immunoreactive bands were detected using the enhanced chemiluminescence detection kit Luminata^TM^Crescendo Western HRP Substrate (Ref: WBLUR0100; Milipore-Sigma, Canada). Data were collected with a ChemiDoc XRS system (Bio-Rad, Canada). The intensities of protein bands were quantified with Adobe Photoshop CS5 software.

### Immunofluorescence

Cells were grown onto 12 or 18mm glass coverslips depending on the experiment. Monolayers were washed twice with PBS 1X and fixed with 4% PFA in PBS for 15 minutes at RT. Cells were permeabilized using 0.3% Triton X-100 in PBS for 10 minutes and subsequently blocked with a 10% skim milk solution in PBS. Alternatively, cells were permeabilized with 0.05% saponin in PBS containing 0.2% BSA for 20 minutes. Cells were incubated with the following antibodies: Purified mouse anti-sorting nexin 1 (Ref: 611482; 1/100, ON 4°C), purified mouse monoclonal anti-sorting nexin 2 (Ref: 611308; 1/100, ON 4°C), mouse monoclonal anti-sorting nexin 6 antibody (D-5) (Ref: sc-365965; 1/100, ON 4°C) customized rabbit polyclonal anti-IBDV VP3 (1/500, 1 hrs 30 min, RT), customized rabbit polyclonal anti-IPNV VP3 (1/800, 1 hrs 30 min, RT), Anti-dsRNA monoclonal antibody J2 (Ref 10010200; 1/50, ON 4°C), mouse monoclonal anti-EEA1(Ref: 61056; 1/100, ON 4°C). After ten washings with PBS 1X monolayers were incubated for 1 hrs 30 min at RT with the corresponding fluorophore-conjugated secondary antibodies described above. Secondary antibody dilutions ranged from 1/500 to 1/1500 depending on the primary antibody. Cells were washed as described above and mounted as described above. Alternatively, cells were incubated with Dapi 1/500 to stain the nucleous, before mounting. For the staining of Lamp-1, cells were fixed/permeabilized with cold methanol at -20°C for 10 min and post-fixed with 4% PFA as described above. Cells were washed twice with PBS 1X and blocked as described above. Cells were incubated with mouse monoclonal anti-Lamp1 (Ref: H4A3; 1/300) for 45 minutes at 37°C, after which the cells were washed, incubated with secondary antibodies and mounted as described above. goat anti-rabbit).

### Opsonization of latex beads and synchronized phagocytosis of IgG-opsonized beads

Polystyrene beads size 2.87um (Ref: PS05002; Bang laboratories, USA) were washed twice with PBS 1X and opsonized with purified human IgG at 4mg/mL in PBS for 2 hrs at RT and washed three more times to remove excess IgG, after which were used for phagocytosis experiments as described below.

To calculate the phagocytic index of QM7 and QM7-VP3 cells, cells were grown as described above. Cells were given opsonized beads and the time course for phagocytosis was synchronized for 15 min and 60 min as described above. External labelling of beads was performed to distinguish internal and external beads anti-Human IgG-Alexa Fluor 647 (1/800) for 1 hr 30 min at RT, after which the cells were washed and mounted as described above.

For PtdIns(3)P kinetics, QM7 and QM7-VP3 cells were grown onto 12 mm glass coverslips in 24-well plates, co-transfected with myc-FcγRIIA and GFP-2FYVE and allowed to reach approximately 80% confluence. Cell monolayers were washed twice with PBS, after which cold DMEM complete cell culture media was added and cells left at 4°C for 10 min. This medium was replaced with a 4°C solution containing the opsonized beads and cells were centrifuged at 300× g for 5 minutes at 4°C in a refrigerated benchtop centrifuge (Eppendorf Centrifuge 5810R, Fisher Scientific, Canada). Working on ice, monolayers were washed with cold PBS 1X until unbound beads were removed and warm media added to allow for the phagocytosis over a time course (10 min, 30 min, 60 min) at 37°C, fixing cells using 4% PFA at each time point and processed as described above. Among beads inside of cells, the fraction positive for GFP-2FYVE was quantified by analyzing cells on Volocity software.

### VP3 interactome analysis

Recombinant His-tagged IBDV VP3, IPNV VP3, and GFP were expressed in *E. coli*, purified chromatographically, and immobilized on magnetic beads. Pre-cleared HeLa cell lysates were incubated with protein-coated beads, and associated proteins were analyzed by either LC-MS/MS or immunoblotting. Complete protein purification, pull-down, and interactome analysis procedures are provided in supplementary information.

### Real time qPCR (RT-qPCR)

CHSE/F cells were cultured at 90% confluency in 25 cm^3^ tissue culture flasks and mock-infected or infected with IPNV at an MOI of 1 PFU/mL. Cells were incubated at RT on an orbital shaker for 1 hr, after which the monolayers were washed with PBS and L-15 complete cell culture media was added. At 2 hrs p.i. the media was replaced with DMSO or 1µM Vps34-IN1 in L-15 complete cell culture media, and the infections were left to proceed for another 18 hrs. Monolayers were washed twice with PBS 1X and processed for RNA extraction. Total RNA was purified with the TRIzol® Plus RNA Purification Kit (Ref:12183555; ThermoFisher Scientific, Canada) by following the manufacturer instructions. RNA concentration and purity were determined with a NanoDrop 1000 and single-use aliquots (20µl) placed at -80°C until use. RT-qPCR analyses were performed via one-step RT-qPCR using the RNA-to-CT™ 1-Step Kit (Ref: 4389986; ThermoFisher Scientific, Canada) and the Applied Biosystems™ QuantStudio™ 3 Real-Time PCR System (ThermoFisher Scientific, Canada). Primers to amplify the IPNV Segment A and IPNV VP3 as well as for the housekeeping gene beta-actin were designed elsewhere(53–55) and purchased via Integrated DNA Technologies (IDT, USA). IPNV Segment A and IPNV VP3 primers were used at the concentration of 100nM, while actin primers were used at 150nM. Samples were run by triplicate, and in three independent trials. Fold change was calculated using the comparative CT method as described in(56).

### Cell viability assay

Cell viability was determined using the MTT viability assay kit (Ref: ab211091; Abcam, Canada) following the manufacturer’s instructions. Briefly, CHSE/F cells were seeded in a 96-well plate and either mock-infected or infected with IPNV at an MOI of 1 PFU/cell. At 2 hr p.i., the cells were treated with DMSO or 1 µM Vps34-IN1 for 18 hrs, after which the medium was discarded, serum-free medium containing MTT reagent was added, and the cells were incubated at 37°C for 3 h. After the addition of MTT solvent, the plate was swirled for 15 min at RT, and the absorbance was measured at an optical density (OD) of 590 nm using a Synergy H4 hybrid microplate reader (BioTek, Canada).

### PolyPIPosomes pull down assays

PolyPIPosomes pull down assays were performed following the manufacturer’s instructions, with some modifications. Briefly, 200 µL of reaction buffer (20 mM HEPES pH 7.4, 120 mM NaCl, 1 mM EGTA, 1 mM MgCl2, 1 mg/mL BSA, 0.2 mM CaCl2, 5 mM KCl) were mixed with ezither 5µL of biotin-coupled control PolyPIPosomes (Ref: Y-0000; Echelon Biosciences, USA) or 5µL biotin-coupled PI3P PolyPIPosomes (Ref: Y-P003; Echelon Biosciences, USA), after which 0.5µg of purified IBDV-VP3 or IPNV-VP3 protein (purified as described above) were added, the samples vortexed for 5 seconds at low speed and incubated for 1h at room temperature using a lab rotator. Afterwards, 20µL of pre-washed MagnaBind™ Streptavidin Beads (Ref: 21344; Thermo Fisher Scientific, Canada) were added to the mix and samples incubated for another 1 h at RT. Beads were pulled down with a magnetic separation rack. Supernatants (inputs) were recovered and the pellets washed three times in 1 mL of reaction buffer and resuspended in 20 µL of reaction buffer. All samples were mixed with 2x Laemmli sample buffer, 4 mM DTT and 1 µM of protease inhibitor and processed by 4–20% Mini-PROTEAN® TGX™ Precast Gel (Ref: 4561093EDU, BioRad, Canada), transferred and immunoblotted as described above.

### Lipid-protein overlay assays

Recombinant His-tagged IBDV VP3, IPNV VP3, GFP, and the PI3P-binding PX-GFP control were expressed in *E. coli* and purified by Ni-NTA affinity chromatography. Phosphoinositides immobilized on nitrocellulose membranes were probed with purified proteins, and binding was detected using anti-His immunoblotting. Detailed protein purification and overlay procedures are provided in supplementary information.

### Fluorescence microscopy and image analysis

Fixed and live-cell imaging was performed using spinning-disk confocal microscopy. Z-stack images and time-lapse sequences were analyzed using Volocity and ImageJ. Vesicle tracking, colocalization, fluorescence intensity measurements, and three-dimensional reconstructions were quantified from multiple cells across independent experiments as detailed in supplementary information.

### On-section Correlative Light Electron Microscopy (CLEM)

QM7 cells stably expressing mKate2-IBDV VP3 were processed by high-pressure freezing, freeze substitution, and Lowicryl HM20 embedding. Thick sections were imaged by fluorescence microscopy and electron tomography. Detailed procedures are provided in SI Appendix.

### Statistical analysis

Unless otherwise specified, data are presented as the mean ± standard deviation (SD) of three independent experiments.

Statistical significance between the control and test groups was determined using the Student’s t-test using KyPlot software version 2.0 (Kyens Lab Inc., Japan). A p-value of < 0.05 was considered to indicate a statistically significant difference.

Alternatively, Chi-square analysis using GraphPad Prism 7.03 was used to determine statistical difference between the distribution of VP3 between two conditions. Values are given as a frequency distribution among cells from three independent trials.

## Author Contributions

Maria Cecilia Gimenez (MCG) and Mauricio Roberto Terebiznik (MRT) are responsible for the overall design of this study. MCG and Diego Ferrero (DF) carried out experiments, data analysis, and interpretation. Mariam Issa (MI), Serene Moussaoui (SM), Liyuan Liu (LL), Suriakarthiga Ganesan (SG) and Micaela Cabrini (MC) carried out experiments and data analysis. Durga Acharya (DA) and Mandy Boermel (MB) carried out CLEM analysis at the EMBL Electron Microscopy Core Facility-Heidelberg. Amanda Rebouças Paixão (ARP) participated in data analysis. MCG and MRT wrote the manuscript. DF, SM, Vanina Zarember (VZ) and Nuria Verdaguer (NV) edited the manuscript. All authors gave approval to the final version of the manuscript.

## Competing Interest Statement

Authors have no competing interests to declare.

## Data, Materials, and Software Availability

All study data are included in the article and/or SI Appendix, except the full proteomics data which is available upon request.

## Acknowledgments

The authors thank the Center for Microscopy and Bioanalysis (CMB) at the University of Toronto Scarborough for the use of the imaging facility. We thank William Lineses and Rachel Chang for technical support and troubleshooting during the experimental setup. The authors thank the EMBL Electron Microscopy Core Facility-Heidelberg for the use of the imaging facility and assistance during preparation and analysis of CLEM samples. The authors thank the Donnelly Mass Spectrometry Center (University of Toronto, Canada), in particular Dr Edyta Marcon, for the processing and bioanalysis of LC-MS/MS samples.

The authors thank Dr. Peter J. Cullen (School of Biochemistry, Faculty of Life Sciences, University of Bristol, UK) for kindly sharing the HeLa KO SNX 5/6; Dr J. Wade Harper (Department of Cell Biology, Blavatnik Institute, Harvard Medical School, USA) for kindly sharing the 293^EL^ cells and Dr. David L. Stachura (Department of Biological Sciences, California State University, USA) for kindly sharing the ZKS cell line. The authors thanks Dr Sergio Grisntein and Dr Michelle Maxson (The Hospital for Sick Children Research Institute, Toronto, ON, Canada) for kindly sharing the recombinant SidK (SidK-AL568). Animations were created using a licensed BioRender account at the name of MCG.

This work was supported by the following parties: Discovery grant from the Natural Sciences and Engineering Research Council to MRT (RGPIN-2024-05569); University of Toronto Scarborough International Research Fund. OVPRI-UTSC to MRT and NV; iNEXT-Discovery (Tech-Sci). Funding for training in microscopy in the Global Bioimaging center at EMBL laboratories Heidelberg, Germany. UTSC VPR-RCF Principal Investigator, collaborative research. MCG was a recipient of a UTSC postdoctoral fellowship. S.M. is a recipient of an NSERC PhD Student Research Award. The CMB facility is funded by a Canada Foundation for Innovation (CFI) grant (#493864). The work at IBMB was funded by the Spanish Ministry of Science and Innovation PID2023-148560NB-I00 to NV. The work at UCalgary was supported by a Discovery Grant from the NSERC to V. Zaremberg.

## Notes

### Competing Interest Statement

The authors have declared no competing interest.

