## Supplementary figures and images for "A dsRNA Virus Converts Early Endosomes into Stable Replication Organelles Through Maturation Arrest and Retrograde Transport"

### Figure S1.tif

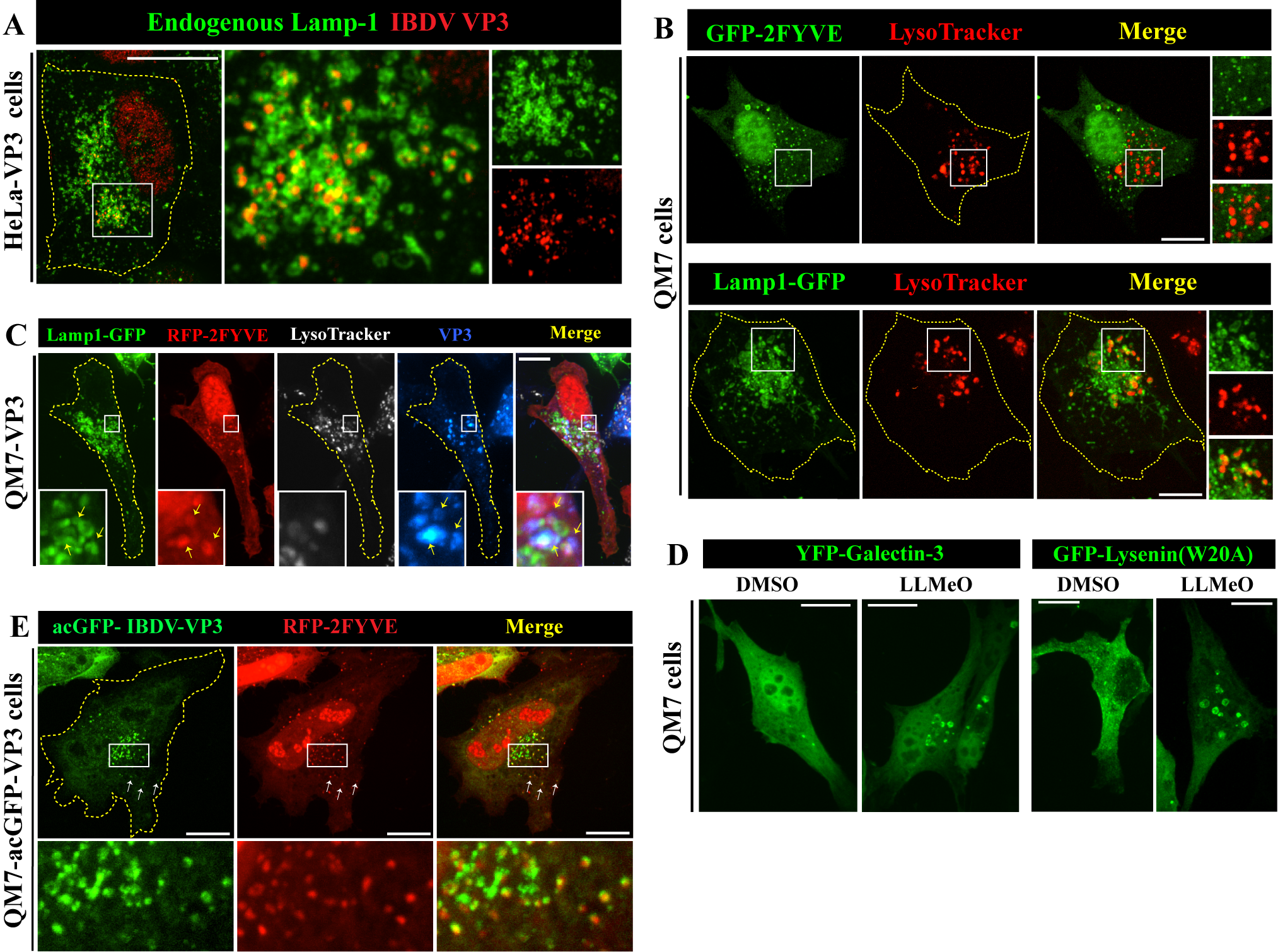

### Figure S2.tif

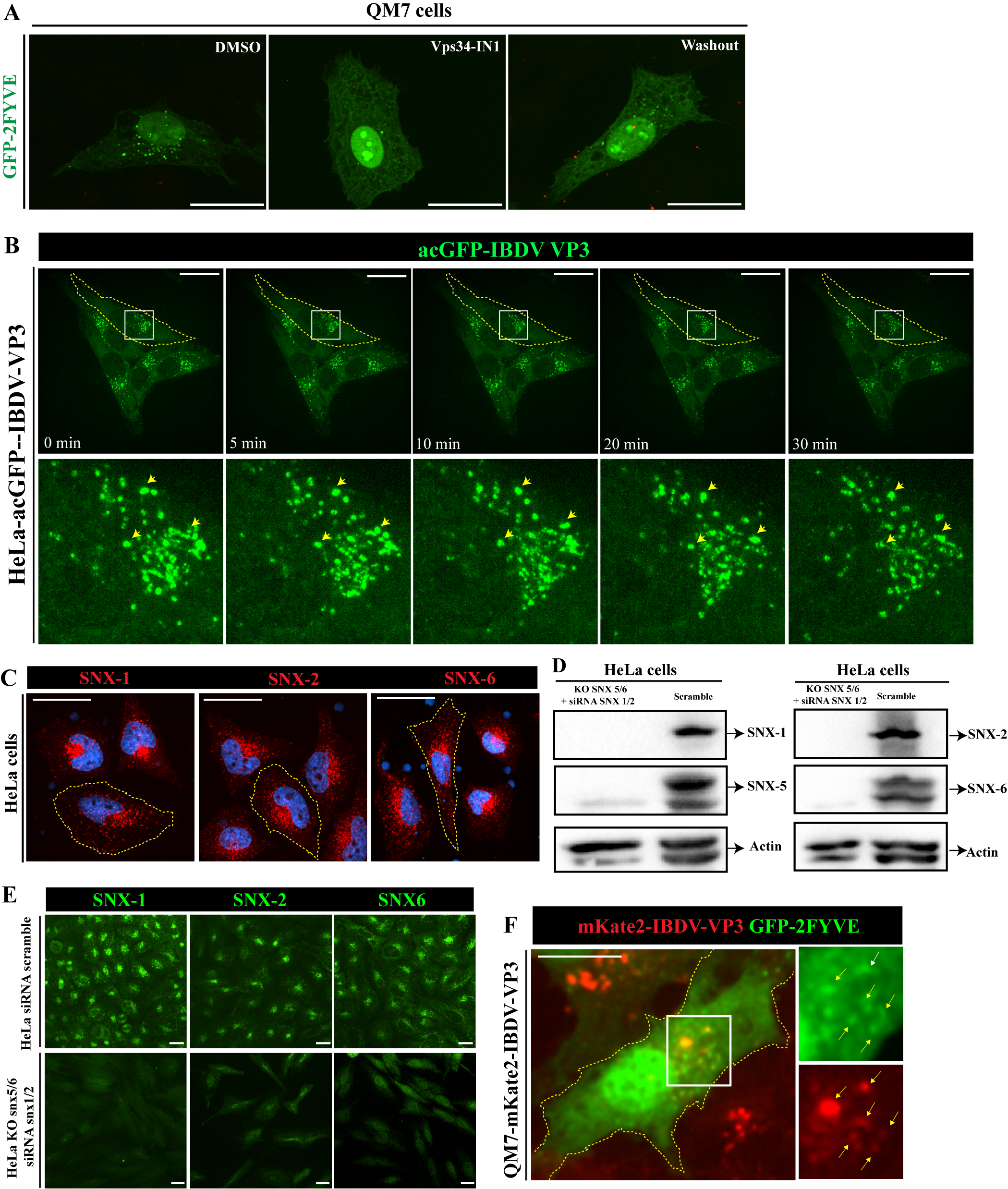

### Figure S3.tif

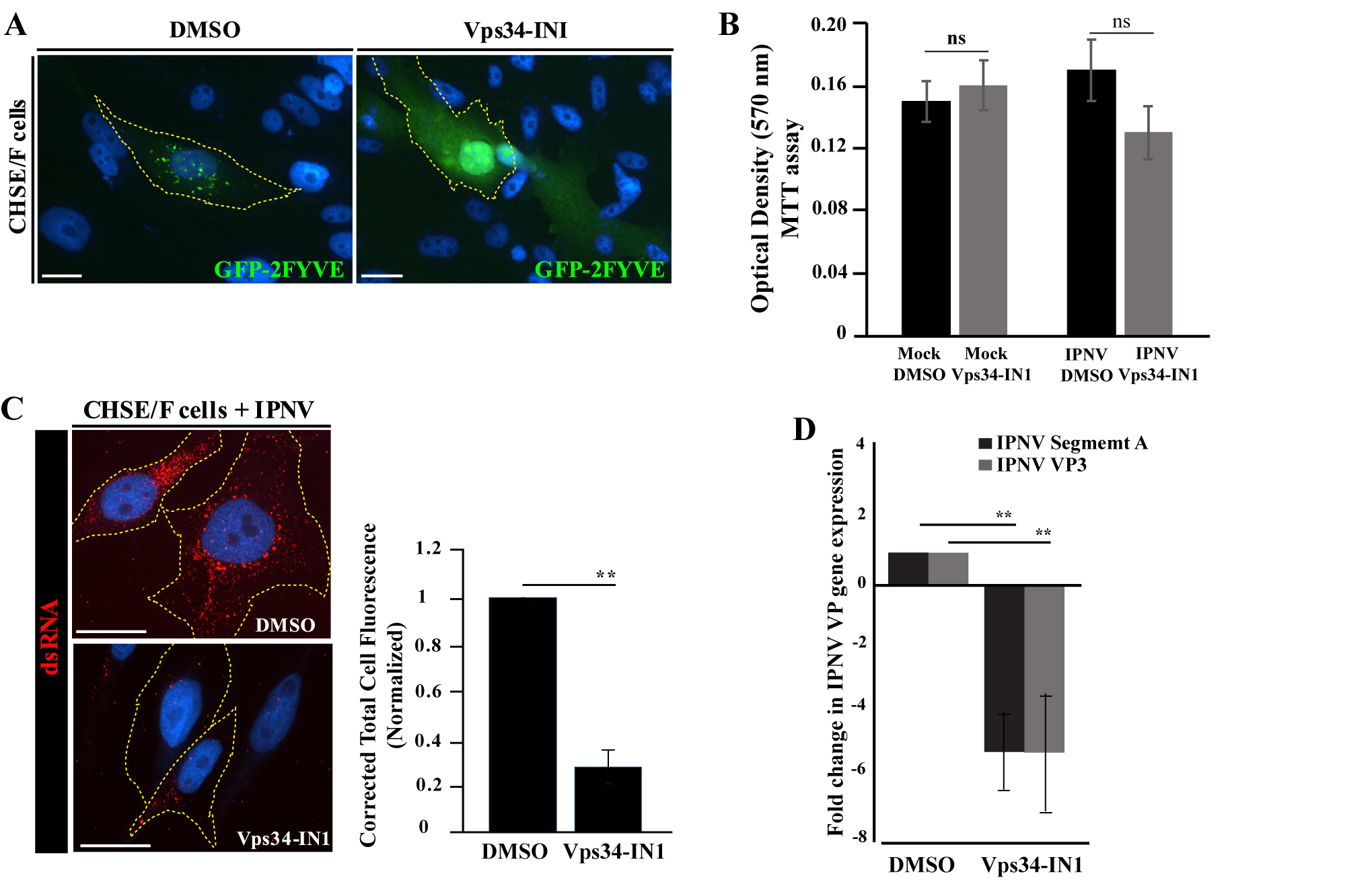

### Figure S4.tif

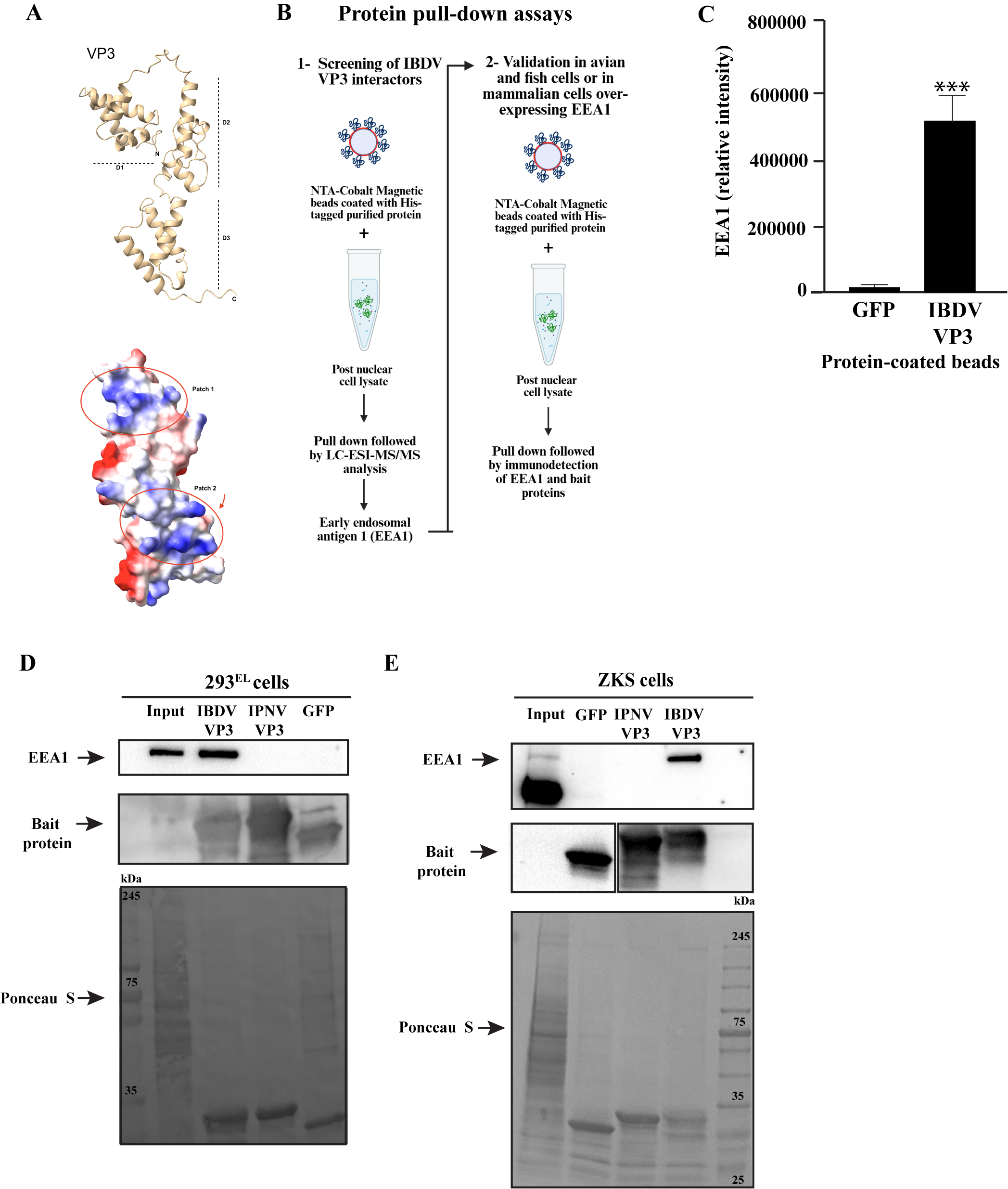
