## Supplementary material for "A dsRNA Virus Converts Early Endosomes into Stable Replication Organelles Through Maturation Arrest and Retrograde Transport": SI appendix

**This PDF file includes:**

Supplementary Materials and Methods

Figures and captions for S1 to S4

Captions for movies S1 to S3

SI References

**Supplementary Materials**

**Cell lines and culture conditions**

Quail muscle myoblast cells (QM7, ATCC CRL-1962) were cultured in Dulbecco’s modified Eagle’s medium (DMEM; Gibco, Thermo Fisher Scientific) supplemented with 10% fetal bovine serum (FBS; Gibco, Thermo Fisher Scientific) and 25 mM HEPES (Gibco, Thermo Fisher Scientific). Stable QM7 cell lines expressing untagged IBDV VP3 (QM7-VP3), mKate2-tagged VP3 (QM7-mKate2-VP3), or acGFP-tagged VP3 (QM7-acGFP-VP3) were maintained in complete medium supplemented with G418 sulfate (Geneticin; Gibco) at 0.5 mg/mL. HeLa cells (ATCC CCL-2) were cultured in DMEM supplemented with 10% FBS. Stable HeLa cell lines expressing untagged VP3 (HeLa-VP3), mKate2-tagged VP3 (HeLa-mKate2-VP3), or acGFP-tagged VP3 (HeLa-acGFP-VP3) were maintained in complete medium containing G418 sulfate at 1 mg/mL. HeLa SNX5/6 KO cells and 293EL cells were cultured as previously described (1, 2). All mammalian and avian cell lines were maintained at 37°C in a humidified incubator with 5% CO₂. CHSE/F cells (formerly CHSE-214; Sigma-Aldrich, Ref. 00021714) were cultured in Leibovitz's L-15 medium (Gibco, Thermo Fisher Scientific) supplemented with 10% FBS at 20°C. For infection experiments, cell cultures were maintained at 14°C. For experiments requiring CHSE/F cell attachment to glass coverslips, coverslips were coated with 6 µg/mL fibronectin in PBS 1X (PBS). Coverslips were completely submerged in coating solution and incubated for 1 hour (hr) at room temperature (RT) or overnight (ON) at 4°C. The solution was then aspirated, and cells were seeded in complete L-15 medium as described. The ZKS cell line was generated and maintained as previously described (3)

**Stable cell lines development**

To generate stable pools of HeLa-VP3, HeLa-mkate2-VP3, HeLa-acGFP-VP3, QM7-mkate2-VP3 or QM7-acGFP-VP3, cells were transfected with the corresponding plasmid and selection proceeded as follows. First, kill curves were established for G418 in both cell lines. Next, cells were seeded at a density of 125 cells/well on dedicated 24-well plates and cultured for 12 hours (hrs), after which cells were transfected with Lipofectamine 3000 (Invitrogen, Thermo Fisher Scientific) following the manufacturer's recommendations. Transfections were performed using a 0.5:1 ratio of plasmidic DNA (µg) to Lipofectamine (µl) for QM7 cells, or a 1:1 ratio for HeLa cells. After 12 hrs, cells were cultured in 1 ml of selection medium containing the corresponding complete media and 1 mg/ml of G418. Selection media were replaced every 3 days during a 21-day period. Surviving pools were expanded according to their growth rate and characterized by fluorescence microscopy.

**Virus propagation and quantification**

The infectious bursal disease virus (IBDV) Soroa strain was propagated in QM7 cells as previously described (4). Briefly, cells at approximately 70% confluency were infected at an MOI of 0.05 PFU/cell. After complete cytopathic effect (72-96 h post-infection), cells and supernatants were collected and subjected to three freeze-thaw cycles consisting of 15 min at −80°C followed by 5 min at 37°C. Cell lysates were clarified by centrifugation at 4°C and 10,000 × g to remove cellular debris, and cleared supernatants were mixed with a PEG precipitation solution (20% PEG 8000 and 3 M NaCl; 1:4 volume ratio) and incubated ON at 4°C. Viruses were pelleted by centrifugation at 4°C and 12,000 × g for 30 minutes (min) and resuspended in PES buffer (25 mM PIPES, pH 6.2, 150 mM NaCl, and 20 mM CaCl₂). Intracellular and extracellular IBDV titers were determined by the plaque assay in QM7 cells as previously described (5).

The infectious pancreatic necrosis virus (IPNV) Sp strain (ATCC VR-1318) was propagated in CHSE/F cells. Cells at approximately 70% confluency were infected at an MOI of 0.05 PFU/cell and incubated on an orbital shaker for 2 hrs at RT to facilitate viral adsorption. Medium was then supplemented to a final concentration of 2% FBS and cultures were maintained at 14°C until complete cytopathic effect was observed (72-96 h post-infection). Cells and supernatants were harvested and processed as described above for IBDV. PEG-concentrated IPNV preparations were further purified by sucrose cushion ultracentrifugation. Discontinuous gradients were prepared in Ultra-Clear centrifuge tubes (Beckman Coulter) by sequentially layering 2 mL of 50% (w/v) sucrose and 2 mL of 25% (w/v) sucrose in PES buffer. PEG-concentrated virus (0.5 mL) was carefully layered onto the gradients and centrifuged at 159,000 × g for 1 h at 4°C using an SW55 Ti swinging-bucket rotor (Beckman Coulter). Viral particles recovered from the interface above the 25% sucrose layer were collected, aliquoted, and stored at −80°C.

Intracellular and extracellular IPNV titers were determined by endpoint dilution (TCID₅₀) assay followed by Reed-Muench analysis. CHSE/F cells were grown in L-15 medium containing 10% FBS and detached at full confluency. Cells were resuspended in L-15 medium supplemented with 7.5% FBS. Serial virus dilutions were prepared in U-bottom plates and transferred to 96-well plates (50 µL/well). CHSE/F cells were then added (100 µL/well), yielding a final FBS concentration of 5%. Mock-infected controls received serum-free L-15 medium. Following a 1 h attachment period at 20 ± 2°C, cultures were incubated at 15°C for 3-5 days. Cells were fixed with 10% formaldehyde for 60 min and stained with crystal violet for 20 min. TCID₅₀/mL values were calculated using the Reed-Muench method and converted to PFU/mL using the conversion factor 0.7.

**DNA constructs and transfection methods**

DNA constructs used in this study for expression in eucaryotic cells: RFP-2FYVE, GFP-2FYVE, GFP-p40-PX, pcDNA3.1 IBDV-VP3 full length, pcDNA3.1 IBDV-VP3ΔCt and pcDNA3.1 IBDV-VP3ΔNt were described elsewhere in(4, 6). GFP-Rab5wt, GFP-Rab7wt, Lamp1-GFP, SialT2-YFP or GalNAcT-YFP were described elsewhere in(7, 8). Lamp1-mCherry, RFP-Rab5, RFP-Rab7, Fc Receptorγ2A-Myc, CD63-GFP were kindly provided by Dr Sergio Grinstein (The Hospital for Sick Children Research Institute, Toronto, ON, Canada). GFP-Rab4 was a gift from John H. Brumel (The Hospital for Sick Children Research Institute, Toronto, ON, Canada). YFP-Galectin3 was kindly provided by Dr. Maria Isabel Colombo (IHEM‐UNCuyo, Argentina). The following plasmids were purchased from Addgene: GFP-Rab9a (Plasmid #169071), EGFP-Rubicon (Plasmid # 21636), GFP-EEA1 (Plasmid #42307). GFP-LyseninW20A was kindly provided by Dr. Jonathan Canton, University of Calgary (Calgary Alberta). pEGFP-Mon1a and pEGFP-Ccz1were purchased from RIKEN BioResource Research Center (Gene Engineering Division, Japan). Arl8B-GFP was kindly provided by Dr. Roberto Botelho (Toronto Metropolitan University, Toronto, ON, Canada).

The following plasmids were developed during this study using the Genscript cloning service: pcDNA3.1 acGFP-IBDV-VP3, pcDNA3.1 mKate2-IBDV-VP3 (both subcloned by modification of Addgene Plasmid #128047) and pcDNA3.1(+) IPNV-VP3.

DNA constructs for expression in procaryotic cells: pACYC-duet-tev-VP3-IBDV full length, pACYC-duet-tev-EGFP, pACYC-duet-tevVP3-IPNV full length were synthesized via GenScript. PX-GFP-V5-His6 and GFP-V5-His6 were provided by Dr Vanina Zaremberg (University of Calgary, Canada).

HeLa and QM7 cells were transfected with Lipofectamine 3000 (Invitrogen, Canada) following the manufacturer’s instructions. For optimizing transfections in CHSE/F cells, cell were transfected in 70% confluent cell monolayers employing 1.5ug of plasmadic DNA and 1.25ul of FugeneHD (Ref: E2311; Promega, Canada) and cells were allowed to express the fluorescent proteins for 24 hours before infections or downstream analysis.

Antibodies and other reagents

The following commercially available antibodies were used in this study: Primary antibodies. Purified mouse anti-sorting nexin 1 (Ref: 611482; BD Transduction Labs, Canada), purified mouse monoclonal anti-sorting nexin 2 (Ref: 611308; BD Transduction Labs, Canada), mouse monoclonal anti-sorting nexin 5 antibody (F-11) (Ref: sc-515215 ; Santa Cruz Biotechnology, USA), mouse monoclonal anti-sorting nexin 6 antibody (D-5) (Ref: sc-365965; Santa Cruz Biotechnology, USA), anti-M6PR monoclonal antibody (IGF2R 2G11) (Ref: MA1-066; ThermoFisher Canada), rabbit polyclonal Anti-α-Actin-1 antibody (Ref: A2066; Millipore Sigma, Canada), Purified mouse anti-GM130 ( Ref: 610822; BD Transduction Labs, Canada), anti-dsRNA monoclonal antibody J2 (Ref 10010200; supplied by Scicons, Hungary), anti -IPNV Vp2 protein mAb (clone 1B3/E10) (Ref: FM130CY-5; Austral Biologicals, USA), mouse anti-IPNV VP3 protein, (clone 6E1/F2) (Catalog No. FM-130DZ-5; Austral Biological, USA), mouse monoclonal anti-Lamp1 (Ref: H4A3; Developmental Studies Hybridoma Bank, USA) mouse monoclonal anti-EEA1(Ref: 61056; BD Transduction Labs); rabbit polyclonal anti-EEA1 (Ref: 2900, Abcam Canada), rabbit polyclonal anti-GFP (Ref: ab290, Abcam Canada), purified human IgG (Ref: I4506; Milipore-Sigma, Canada). Polyclonal serums against IBDV-VP3, IPNV-VP3 and IPNV virus were purchased through Custom Antibody Services to Cederlane Laboratories, Canada. Our laboratory provided the antigens in the form of purified proteins or whole inactivate viral particles, respectively. Secondary antibodies: donkey anti-abbit IgG Alexa Fluor 647 (Ref: A31573; Thermofisher Canada), donkey anti-rabbit IgG Alexa Fluor 555 (Ref: A31572; Thermofisher Canada), oat anti-rabbit IgG Alexa Fluor 405 (Ref: A31556; Thermofisher Canada), donkey anti-rabbit IgG Alexa Fluor 488 (Ref: A21206; Thermofisher Canada), donkey anti-mouse IgG Alexa Fluor 555 (Ref: A31570; Thermofisher Canada), donkey anti-mouse IgG Alexa Fluor 647 (Ref: A315771; Thermofisher Canada), goat anti-mouse IgG Alexa Fluor 488 (Ref: A11001; Thermofisher Canada), goat anti-human IgG Alexa Fluor 647 (Ref: A21445; Thermofisher Scientific, Canada), horseradish peroxidase (HRP)-conjugated goat anti-rabbit immunoglobulin (Ref: A0545; Jackson Immunoresearch, USA), and goat anti-mouse IgG–HRP (Ref: A9044; Jackson Immunoresearch, USA).

Other reagents used in this study: LysoTracker Red DND-99 (Thermo Fisher Scientific, Ref: L7528), recombinant SidK conjugated with Alexa Fluor 568 (SidK-AL568) was kindly provided by Dr Sergio Grisntein (The Hospital for Sick Children Research Institute, Toronto, ON, Canada). The lysosomotropic agent L-leucyl-L-leucine methyl ester (LLOMe, Ref: 4000725, Becham, USA), Brefeldin A (Ref: 20350-15-6), Golgicide A (Ref: G0923) and Nocodazole (Ref: M1404) were purchased from Sigma Milipore, Canada. Vps34-IN1 (Ref: S7980; Selleckchem and Dynarrestin (Ref: E1172) were purchased from Sellechchem, Canada.

**Labelling of acidic compartments**

Cells were seeded onto 12 mm glass coverslips and transfected with either Lamp-1 GFP or GFP-2FYVE using Lipofectamine 3000 reagent. At 20 h post transfection, transfected cells were incubated with 1 μM LysoTracker Red DND 99 diluted with cell culture media for 1 h at 37˚C with 5% CO2. Coverslips were then washed with 1X PBS, fixed with 4% PFA and immunostained against IBDV VP3.

**Detection of vacuolar H^+^ATPase using the** **SidK-AF568 probe**

QM7-VP3 cells were seeded onto 12 mm glass coverslips, transfected with GFP-2FYVE, and fixed with 3% PFA for 10 min at RT. The cells were permeabilized with 0.05% saponin in PBS containing 0.2% Bovine Serum Albumin (BSA) for 30 min at RT and then incubated with 10 μg/mL of SidK-AF568 diluted in saponin for 45 min at RT. The monolayers were washed five times with 1X PBS. Imaging was performed the following day using live-cell imaging chambers with the cells maintained in 1X PBS.

**LLOMe-triggered membrane damage**

QM7 cells were seeded onto 12 mm glass coverslips and transfected with either YFP Galectin-3 or GFP-Lysenin (W20A). At 20 h post transfection, cells were incubated with 200 μM of LLOMe or Dimethyl Sulfoxide (DMSO; BioShop, Catalogue No. DMS666.100) in DMEM for 1 h before being washed with 1X PBS, fixed with 4% PFA and mounted in Fluorescence Mounting Medium Dako (Ref: S3023 Agilent, Canada). QM7 VP3 cells seeded onto 12 mm glass coverslips were co-transfected with either fluorescently tagged endocytic marker GFP-2FYVE/RFP-2FYVE or Lamp1-GFP/Lamp1-mCherry and with either YFP Galectin-3 or GFP-Lysenin(W20A) using Lipofectamine 3000 reagent. At 20 h post transfection, cells were incubated in 200μM of LLOMe or DMSO in DMEM for 1 h before being washed with 1X PBS, fixed with 4% PFA and processed for immunofluorescence against IBDV VP3 and analyzed by spinning disk confocal microscopy as described below.

**siRNA treatment and IBDV infection**

HeLa cells were seeded in 6-well plates and grown to approximately 80% confluency prior to transfection. Cells were transfected with either a non-targeting control siRNA (Silencer Select Negative Control), siRNA targeting **SNX1**, or siRNA targeting **SNX2** (Thermo Fisher Scientific) using Lipofectamine RNAiMAX according to the manufacturer's instructions.

At 48 h post-transfection, culture medium was removed and cells were infected with IBDV at an MOI of 1 PFU/cell. Following a 1 h adsorption period, the inoculum was removed, monolayers were washed once with PBS, and fresh transfection medium was returned to the cells. Infection was allowed to proceed for an additional 24 h.

Following infection, one set of samples was processed for immunoblot analysis, while parallel samples were collected for determination of intracellular and extracellular viral titers as described below.

**Treatment with pharmacological inhibitors**

**VPS34 inhibition.** The activity of VPS34-IN1 against fish VPS34 and its lack of cytotoxicity in CHSE/F cells were validated prior to experimentation (Fig. S3D,E).

For washout experiments, QM7 cells expressing either IBDV VP3 or acGFP-IBDV VP3 were treated with 1 µM VPS34-IN1 in complete medium for 1 h. Cells were washed with PBS and incubated in fresh medium for 10 min to 1 h before analysis.

To assess VPS34 inhibition in fish cells, CHSE/F cells were transfected with GFP-2FYVE and treated with either DMSO or 1 µM VPS34-IN1 for 1 h. Cells were then washed with PBS, fixed in 4% paraformaldehyde, and processed for immunofluorescence as described below.

For infectivity assays, CHSE/F cells grown to approximately 70% confluency in 6-well plates were infected with IPNV at an MOI of 1 PFU/cell for 1 h at RT with orbital shaking. Following virus adsorption, cells were incubated in complete L-15 medium for 2 h at 15°C and subsequently treated with either DMSO or 1 µM VPS34-IN1 for 18 h. Culture supernatants were collected as the extracellular virus fraction. Adherent cells were harvested, subjected to three freeze-thaw cycles, and centrifuged at 16,000 × g for 10 min to remove debris. Intracellular and extracellular fractions were quantified by endpoint dilution assay as described below.

For immunoblot analyses, CHSE/F cells were infected and treated as above, after which cell lysates were prepared for SDS-PAGE and detection of IPNV VP2 and VP3 proteins.

**Golgi and microtubule disruption.** QM7 cells stably expressing mKate2-IBDV VP3 were seeded on 18-mm glass coverslips and treated for 2 h with one of the following inhibitors: Golgicide A (30 µM), Brefeldin A (5 µg/mL), Nocodazole (20 µM), or Dynarrestin (150 µM). Cells were subsequently fixed and processed for fluorescence microscopy as described below.

**Identification of VP3 interactors using protein-coated magnetic beads**

Protein expression and purification. Escherichia coli BL21 (DE3) cells were chemically transformed with pACYC-duet constructs encoding IBDV VP3, IPNV VP3 or GFP. Isolated bacterial colonies were inoculated in Luria Broth (LB) medium supplemented with 50 µg/mL chloramphenicol at 37 °C with shaking at 200 rpm. When the culture reached an optical density at 600 nm (OD₆₀₀) of 0.7–0.8, protein expression was induced by adding 0.1 mM isopropyl β-D-1-galactopyranoside (IPTG). The cultures were then incubated for 18 h at 17 °C. Cells were harvested by centrifugation (JLA-9.1000 rotor, 3000 g), washed with 100 mL of phosphate-buffered saline (PBS), and stored at −20 °C. For protein purification, cell pellets were thawed on ice and resuspended in lysis buffer containing 50 mM Tris-HCl pH 8.0, 500 mM NaCl, 10% glycerol, 1 mM DTT, 10 µg/mL lysozyme, 10 µg/mL DNase (Bio Basic, Canada), 10 µg/mL RNase (Neo-Biotech), and 4 mM MgCl₂, supplemented with a protease inhibitor cocktail (Complete EDTA-free, Roche). Cell lysis was performed using a high-pressure cell disruptor (Constant Systems, UK) at 1.4 kbar and 4 °C. The lysate was clarified by centrifugation at 48,000 × g for 30 min using a JA25.50 rotor (Beckman Coulter). The resulting supernatant was filtered through a 0.45 µm membrane and applied to a HisTrap column (GE Healthcare) using an ÄKTA FPLC system. For affinity purification, 5 mM imidazole was included in the binding buffer. The column was washed with 10 column volumes (CV) of buffer A (50 mM Tris-HCl pH 8.0, 500 mM NaCl, 10% glycerol, 1 mM DTT), followed by elution with a 30 CV linear gradient up to 100% buffer B (buffer A supplemented with 500 mM imidazole). Fractions containing VP3 were pooled, diluted fivefold with 50 mM Tris-HCl (pH 8.0), 10% glycerol, and 1 mM DTT, and subsequently loaded onto a HiTrap Q column (Cytiva). The column was washed with buffer containing 50 mM Tris-HCl pH 8.0, 100 mM NaCl, 10% glycerol, and 1 mM DTT. Proteins were eluted using a 20 CV linear gradient to 1 M NaCl in the same buffer. VP3-containing fractions were pooled and dialyzed against buffer A. A final purification step was performed by size-exclusion chromatography (SEC) using a Superdex 200 10/300 column (GE Healthcare) equilibrated in buffer A supplemented with 2 mM DTT. Fractions were analyzed by SDS-PAGE, and those with the highest purity were pooled. The protein was concentrated using a Centricon filter device (3 kDa molecular weight cutoff, Millipore) and stored at −80 °C.

Sample preparation for pulldown assays: Coating of magnetic beads with His-IBDV-VP3, His-IPNV-VP3 and His-EGFP was performed using Dynabeads™ His-tag isolation and pulldown magnetic beads (Ref: 10103D; Thermo Fisher Scientific, Canada). To saturate the beads, 25 µL of Dynabeads were incubated with 80 µg of purified protein in 1X Binding/washing buffer (50 mM S

odium Phosphate pH 8.0, 300 mM NaCl, 0.01% Tween™-20) for 10 minutes at RT. Beads were washed four times in 1X Binding/washing buffer and coating efficiency was verified via SDS-PAGE followed by Coomassie staining, and western blot analysis using anti-VP3 and anti-GFP primary antibodies and as described above. Cell lysates were prepared and pre-cleared as follows: HeLa cells were cultured in 150 cm² flasks, harvested by trypsinization, and washed twice with PBS 1X. Cell pellets were resuspended in 600 µl NP-40 lysis buffer (150 mM NaCl, 1.0% NP-40, 50 mM Tris-HCl pH 8.0) supplemented with 1% protease and phosphatase inhibitor cocktail (Ref: PPC1010; Sigma-Millipore). After a 10-minute incubation on ice, lysates were centrifuged at 1,000 x g for 5 minutes at 4°C to obtain post-nuclear supernatants (PNS). To minimize non-specific binding, the PNS was pre-cleared by incubation with non-coated magnetic beads for 30 minutes at RT before the pull-down assay.

Pull-down assays for Liquid Chromatography and Mass Spectrometry analysis (LC-MS/MS): Pre-cleared post-nuclear cell lysates (250 µL) were diluted with 450 µL of 1X Pull-down buffer (3.25 mM Sodium Phosphate pH 7.4, 70 mM NaCl, 0.01% Tween™-20) and incubated with 25 µL of VP3- or GFP-coated beads for 30 minutes at RT. Following incubation, beads were washed four times with 1X Binding/washing buffer and three times with a low-salt washing buffer (10 mM Tris-HCl pH 8, 100 mM NaCl). Proteins were eluted from the beads using 0.5 M ammonium hydroxide (Ref: A669500, Fisher Scientific, Canada). A 5 µL aliquot of the eluate was reserved for quality control via silver staining (Pierce™ Silver Stain for Mass Spectrometry, 24600), while the remaining eluates were flash-frozen in liquid nitrogen and stored at -80°C for subsequent processing and bioinformatic analysis at the Donnelly Mass Spectrometry Center (University of Toronto, Canada).

Pull-down assays for western blotting: Pre-cleared post-nuclear cell lysates (250 µL) were diluted with 450 µL of 1X Pull-down buffer (3.25 mM Sodium Phosphate pH 7.4, 70 mM NaCl, 0.01% Tween™-20) and incubated with 25 µL of VP3- or GFP-coated beads for 30 minutes at RT. Following incubation, beads were washed four times with 1X Binding/washing buffer and three times with a low-salt washing buffer (10 mM Tris-HCl pH 8, 100 mM NaCl), after which magnetic beads were resuspended in 50 µL of NP-40 lysis buffer containing 1X Laemmli sample buffer and 2 mM DTT. Samples were boiled at 100°C for 10 minutes, and the supernatant was isolated using a magnet. Protein extracts (30 µL) were resolved by SDS-PAGE using 4–20% Mini-PROTEAN® TGX™ Precast Gel (Ref: 4561093EDU, BioRad, Canada) and proteins transferred as described above. Membranes were washed and immersed in a 0.1% (w/v) Ponceau S solution (in 5% acetic acid) for 10 minutes at RT, after which membranes were rinsed three times with deionized water and two times with PBS 1X and blocked with 10% skim milk in PBS 1X ON at 4°C. Membranes were washed twice with PBS 1X prior to incubation with the following antibodies: customized rabbit polyclonal anti-IBDV VP3 (1/3000), customized rabbit polyclonal anti-IPNV VP3 (1/2000), mouse monoclonal anti-EEA1(1/1000); rabbit polyclonal anti-EEA1 (1/500) and rabbit polyclonal anti-GFP (1/1000). After three washings with PBS–0.05% Tween 20, for 15 min each, membranes were incubated with the corresponding HRP-conjugated secondary antibodies described above. Membranes were washed and the corresponding immunoreactive bands were detected as described above.

**Lipid-protein overlay assay**

Protein production and purification: *Escherichia coli* BL21 cells were chemically transformed with constructs encoding the full length IPNV VP3-Hisx6, full length IBDV VP3-Hisx6, GFP-V5-Hisx6 or PX-GFP-V5-His6. Isolated bacterial colonies were inoculated in 40 mL of LB medium supplemented with 100 µg/mL ampicillin and grown ON at 37 °C. The cultures were then diluted into 800 mL of LB medium, and protein expression was induced with 1 mM IPTG for 4 h at 37 °C. Bacterial pellets were resuspended in 10 mL of lysis buffer [50 mM Tris-HCl pH=8; 500 mM NaCl; 2mM DTT; 0.1% NP-40; 10 ug/mL Lysozyme; 10 ug/mL DNAse; 10 ug/mL RNAse; 4 mM MgCl2; complete protease inhibitors 1X (Roche Diagnostics); 3 ug/mL pepstatin; 1 mM phenylmethylsulfonyl fluoride] and sonicated for 30 sec, 5 times. Cleared lysates were incubated with pre-equilibrated Ni-NTA agarose beads (Qiagen) for 1 h at 4 °C. The slurry was transferred to a chromatography column, washed four times with Buffer A [50 mM Tris-HCl (pH 8.0), 500 mM NaCl, and 15 mM imidazole], and eluted with Buffer B [50 mM Tris-HCl (pH 8.0), 500 mM NaCl, and 250 mM imidazole]. Protein eluates were initially concentrated using Amicon Ultra-15 centrifugal filters (MilliporeSigma). To ensure proper folding, all proteins (except PX-GFP-V5-His6) underwent an on-column refolding procedure. Concentrated proteins were resuspended in Buffer A supplemented with 3.5 M urea, re-bound to the Ni-NTA column, and washed sequentially with step-down gradients of urea (1.8 M, 0.9 M, and 0 M) in Buffer A. The refolded proteins were eluted with Buffer B, concentrated using Amicon Ultra-15 filters, identified by Western blotting, and stored at −80 °C

Preparation of spotted membranes: For lipid-protein overlay assays using a panel of PtdIns species, 200 pmol of each phosphoinositide specie {DOPI (18:1 PI) Avanti 850149-100 ug; DOPI(3)P [18:1 PI(3)P] Avanti 850150-100 ug; DOPI(4)P [18:1 PI(4)P] Avanti 850151-100 ug; DOPI(5)P [18:1 PI(5)P] Avanti 850152-100 ug; DOPI(3,5)P [18:1 PI(3,5)P_2_] Avanti 850154-100 ug; DOPI(4,5)P [18:1 PI(4,5)P_2_] Avanti 850155-100 ug; DOPI(3,4,5)P [18:1 PI(3,4,5)P_3_] Avanti 850156-100 ug} was prepared in a mixture of chloroform:methanol:H2O (20:9:1 v/v) and spotted onto 0.2 µm nitrocellulose membranes (Cat#10600054, GE Healthcare). For concentration-dependent overlay assays, DOPI(3)P was spotted at 50, 100, 200 or 400 pmol per spot using the same solvent system.

Overlay protocol: Spotted membranes were blocked with Tris-buffered saline containing 0.1% Tween-20 (TBST) 1X supplemented with 3mM CaCl2 and 3% fatty acid-free bovine serum albumin (FF-BSA). The membranes were then incubated with 10µg/ml of the purified target protein diluted in the blocking buffer. After washing three times with TBST 1X containing 3mM CaCl2, tagged proteins were detected using an anti-His6 primary antibody (1:2000) followed by horseradish peroxidase (HRP)-conjugated secondary antibody (G21040; Thermo Fisher). Proteins were visualized via chemiluminescence and captured using an iBright FL1500 imaging system (Invitrogen).

**Fluorescence microscopy and image analysis**

Confocal imaging of fixed cells was performed employing Spinning Disc Confocal Microscopes. A Quorum Wave FX-X1 spinning-disk confocal microscope system (Quorum Technologies, Inc.; Canada) equipped with an inverted fluorescence microscope (DMI6000B; Leica Microsystems) using either a Hamamatsu ORCA-R2 camera (C10600-10B) or a Hamamatsu ImagEM Enhanced EM charge-coupled device (CCD) camera (Hamamatsu Corporation) and operated via MetaMorph software. Images were acquired using either 40× oil immersion objectives (1.3 NA) or 63× oil immersion objectives (1.4 NA). Alternatively, a Leica DM16000B spinning-disk confocal microscope (Quorum Technologies, Inc.; Canada) equipped with an ORCA-R2 camera (Hamamatsu, Japan) also operated via MetaMorph software was used to acquire images, using a 63× oil immersion objectives (1.4 NA). Unless otherwise specified, all images are presented as the merge of Z-stacks.

The generation of a green, fluorescent VP3 construct, created by tagging IBDV-VP3 with the monomeric *Aequorea coerulescens* GFP (acGFP -VP3) (9) enabled us to acquire live cell imaging of VP3 dynamics. To this end, 18mm glass coverslips were enclosed in a Leiden chamber and immersed in Live Cell Imaging Solution from Molecular Probes™ (Ref: A59688DJ; Thermo Fisher, Canada) supplemented with 10% FBS and mounted in a microscope-mounted environmental chamber maintained at 37°C. Time-lapse videos were acquired using the Leica DM16000B spinning-disk confocal microscope and a 63X oil immersion objective lens. Image and time-lapse video processing, and analysis were performed with Volocity 6.1 (Volocity®, PerkinElmer Inc. Canada) and/or ImageJ software (10).

The fluorescence intensity profiles and 3D rendering were generated in Volocity®.

Vesicle tracking was performed manually from timelapse confocal images using the plugin Manual Tracking (ImageJ software). A total of 5 cells were analyzed, and for each cell, at least 10 individual vesicles at the periphery of the cell or at the perinuclear area were tracked. Vesicles coordinates (x,y) were postprocessed in R programming software.

To reconstruct vesicles trajectory, vesicles were tracked individually with the plugin Manual track on Image J software. 5 vesicles were tracked for each cell, and the trajectories (x,y) were originated simultaneously with the tool Overlay dots and lines for all the stacks, which draws the path of the object.

The Volocity Measurements program was utilized to determine the percentage of VP3 objects positive for 1 or 2 endocytic markers. At least 20 cells were analyzed per condition, per experiment.

The corrected total cell fluorescence (CTCF) was calculated using ImagJ and the following formula: CTCF = integrated density – (area of selected cell x means fluorescence of background) and values were normalized.

**On-section Correlative Light Electron Microscopy (CLEM)**

CLEM analysis was done in collaboration with the Electron Microscopy Core Facility EMBL Heidelberg, Germany and as previously described(11, 12). Briefly, QM7 cells stably expressing mKate2-IBDV-VP3 were grown on 0.05-mm-thick, 3-mm- diameter sapphire disks (Ref: 405; Wohlwend GmbH, Switzerland) in DMEM complete cell culture media. The sapphire disks were carbon-coated, and an asymmetric mark etched into the carbon film; before burning the branding in ON at 130°C. Cells grew best on the non-carbon-coated side of the sapphires. Sapphire disks with cells were high-pressure frozen in 3-mm-wide, 40-µm-deep aluminum planchettes (Wohlwend GmbH, Switzerland) using a HPM010 from AbraFluid (Switzerland). The samples were freeze-substituted using a EM-AFS2 (Leica Microsystems, Germany) with 0.1 % Uranyl Acetate (UA) in acetone at -90°C for 24 hrs. The temperature was then raised to -45°C at 5°C/hr and samples were further incubated for 5 hrs. After rinsing in acetone, the samples were gradually infiltrated in Lowicryl HM20 resin, while raising the temperature to -2°C and left to polymerize under UV light for 48 hrs at -25 °C and for further 9 hrs while the temperature was gradually raised to 20°C (5 °C/h). Thick sections (300 nm) were cut from the polymerized resin block and picked up on carbon coated mesh grids. The fluorescence microscopy imaging of the sections was carried out as previously described(13) using a widefield fluorescence microscope (Olympus IX81 with MT20 illumination system equipped with an Olympus PlanApo 100X 1.40 NA oil immersion objective. 10 nm protein A gold (UMC Utrecht, Cell Microscopy Core) was added to the sections as (tomographic) fiducial markers, before post-staining with UA and lead citrate. Tilt series of the areas of interest were acquired over a -60° to +60° tilt range (1° increment) using a FEI Tecnai F30 TEM (300kV, Gatan OneView camera; final pixel size 1.55 nm) and tomograms were reconstructed using the software package IMOD. Correlation between light and electron micrographs was carried out with the plugin ec-CLEM(14) of the software platform Icy. The coordinates of pairs of corresponding features in the two imaging modalities were used to calculate a similarity transform (translation/rotation/scaling), to map the coordinates of the fluorescent spots of interest and to overlay them on the electron tomogram.

Supplementary Figures and Movies

**Supplementary Figure 1.** (A) SDCM images of HeLa-VP3 cells stained for VP3 (red) and Lamp1 (green). Magnified views show representative VP3-Es. (B) SDCM images of QM7 cells expressing GFP-2FYVE or Lamp1-GFP (green) and labeled with LysoTracker (red). Magnified views show representative endocytic compartments. (C) SDCM images of QM7 cells expressing Lamp1-GFP (green) and RFP-2FYVE (red) labeled with LysoTraker (gray) and stained for VP3 (blue). Magnified views show representative VP3-Es. Yellow arrows indicate individual VP3-Es. (D) SDCM images of QM7 cells expressing YFP-galectin-3 or GFP-lysenin W20A (green) following LLOMe treatment. (E) SDCM images of QM7-acGFP-IBDV-VP3 cells expressing RFP-2FYVE (red). White arrows indicate peripheral VP3/PtdIns(3)P-positive compartments; magnified views show representative perinuclear VP3/PtdIns(3)P-positive compartments. Images are representative of three independent experiments. Scale bars, 10 μm.

**Supplementary Figure 2.** (A) SDCM images of QM7 cells expressing GFP-2FYVE (green) following treatment with Vps34-IN1 and drug washout. (B) Representative frames from time-lapse video microscopy of HeLa-acGFP-IBDV-VP3 cells. Magnified views show representative VP3-Es. Yellow arrows track individual VP3-Es over time in the perinuclear region. (C) SDCM images of HeLa cells stained for SNX1, SNX2, and SNX6. (D) Immunoblot analysis of HeLa cells depleted of SNX1/2/5/6. (E) SDCM images of HeLa cells depleted of SNX1/2/6. (F) SDCM images of QM7-mKate2-IBDV-VP3 cells (red) expressing GFP-2FYVE (green). Magnified views show VP3-Es. Yellow arrows indicate VP3/PtdIns(3)P-positive compartments in the perinuclear region. Images and immunoblots are representative of three independent experiments. Scale bars, 10 μm.

**Supplementary Figure 3.** (A) SDCM images of CHSE/F cells expressing GFP-2FYVE (green) and treated with Vps34-IN1. Nuclei are shown in blue. (B) Cell viability analysis of Vps34-IN1-treated CHSE/F cells measured by MTT assay. Values represent mean ± SD from three independent experiments (ns, not significant). (C) SDCM images of Vps34-IN1-treated CHSE/F cells infected with IPNV (MOI = 5) and stained for dsRNA at 8 h postinfection. Quantification of corrected total cellular fluorescence (CTCF) is shown as mean ± SD from three independent experiments (**P < 0.05). (D) RT-qPCR analysis of CHSE/F cells treated with Vps34-IN1 and infected with IPNV. Fold changes for Segment A and VP3 transcripts are shown as mean ± SD from three independent experiments (**P < 0.05). Images are representative of three independent experiments. Scale bars, 10 μm.

**Supplementary Figure 4.** (A) AlphaFold3-predicted structure of full-length IPNV VP3 (residues 1-237). Protein domains (D1-D3) are indicated. Surface electrostatic potential of the D2-D3 core is shown from negative (red) to positive (blue) charge. Predicted PATCH1 and PATCH2 regions are highlighted. (B) Schematic workflow of VP3 pull-down assays followed by LC-ESI-MS/MS or immunoblot analysis. Created in BioRender by MCG (C) Relative abundance of EEA1 in GFP and IBDV-VP3 pull-down samples. Values represent mean ± SD from three independent experiments (***P < 0.01). (D) Pull-down assays using protein-coated magnetic beads and lysates from 293EL cells. GFP served as a heterologous control. (E) Pull-down assays using protein-coated magnetic beads and lysates from ZKS cells. GFP served as a heterologous control. Graphs represent mean ± SD as indicated. Blots are representative of two independent experiments.

**Supplementary Video 1.** Time-lapse video microscopy of QM7-acGFP-IBDV-VP3 captured using SDCM. Frame rate 4 fps.

**Supplementary Video 2.** Time-lapse video microscopy of QM7-acGFP-IBDV-VP3 captured using SDCM. QM7-acGFP-IBDV-VP3 cells were treated with Vps34-IN1 as described in Materials and Methods. Frame rate 4 fps.

**Supplementary Video 3.** Time-lapse video microscopy of HeLa-acGFP-IBDV-VP3 captured using SDCM. Frame rate 4 fps.
